# PERK orchestrates MERCS formation and mitochondrial remodelling promoting physiological adaptations during adaptive UPR signalling

**DOI:** 10.1101/2025.01.20.633960

**Authors:** Jose C. Casas-Martinez, Qin Xia, Penglin Li, Maria Borja-Gonzalez, Antonio Miranda-Vizuete, Emma McDermott, Peter Dockery, Leo R. Quinlan, Katarzyna Goljanek-Whysall, Afshin Samali, Brian McDonagh

**Affiliations:** Discipline of Physiology, School of Pharmacy and Medical Sciences, University of Galway, Ireland; Apoptosis Research Centre, University of Galway, Ireland; Galway RNA Research Cluster, University of Galway, Ireland; Department of Orthopedics, Tongji Hospital, Tongji Medical College, Huazhong University of Science and Technology, China; School of Biological and Chemical Sciences, University of Galway, Spain; Instituto de Biomedicina de Sevilla, IBiS/Hospital Universitario Virgen del Rocío/CSIC/Universidad de Sevilla, Spain; Centre for Microscopy and Imaging, Discipline of Anatomy, University of Galway, UK; Institute of Life Course and Medical Sciences, University of Liverpool, UK

**Keywords:** ER stress, mitochondrial dynamics, ageing, mitochondrial-ER contact sites, myogenesis, *C*. *elegans*

## Abstract

The transfer of information and metabolites between mitochondria and the ER is mediated by mitochondria-ER contact sites (MERCS), facilitating adaptations following changes in cellular homeostasis. MERCS are dynamic structures, essential for maintaining cellular homeostasis through modulation of calcium transfer, redox signalling, lipid transfer, autophagy and mitochondrial dynamics. Acute ER stress in myoblasts promoted myogenesis that required the PERK arm of the UPR^ER^ for increased MERCS assembly, mitochondrial turnover and function. Similarly, induction of acute UPR^ER^ during early development in *C. elegans* resulted in increased lifespan and healthspan. Adaptive UPR^ER^ signalling in myoblasts and *C. elegans*, increased MERCS assembly and activated autophagy, ultimately promoting mitochondrial remodelling. Adaptations were dependent on the developmental stage, as treatment of myotubes or adult *C. elegans* resulted in a maladaptive response. The results identify that PERK is required for increased mitochondrial ER communication in response to adaptive UPR signalling, promoting mitochondrial remodelling and improved physiological function.

**Graphical Abstract:** 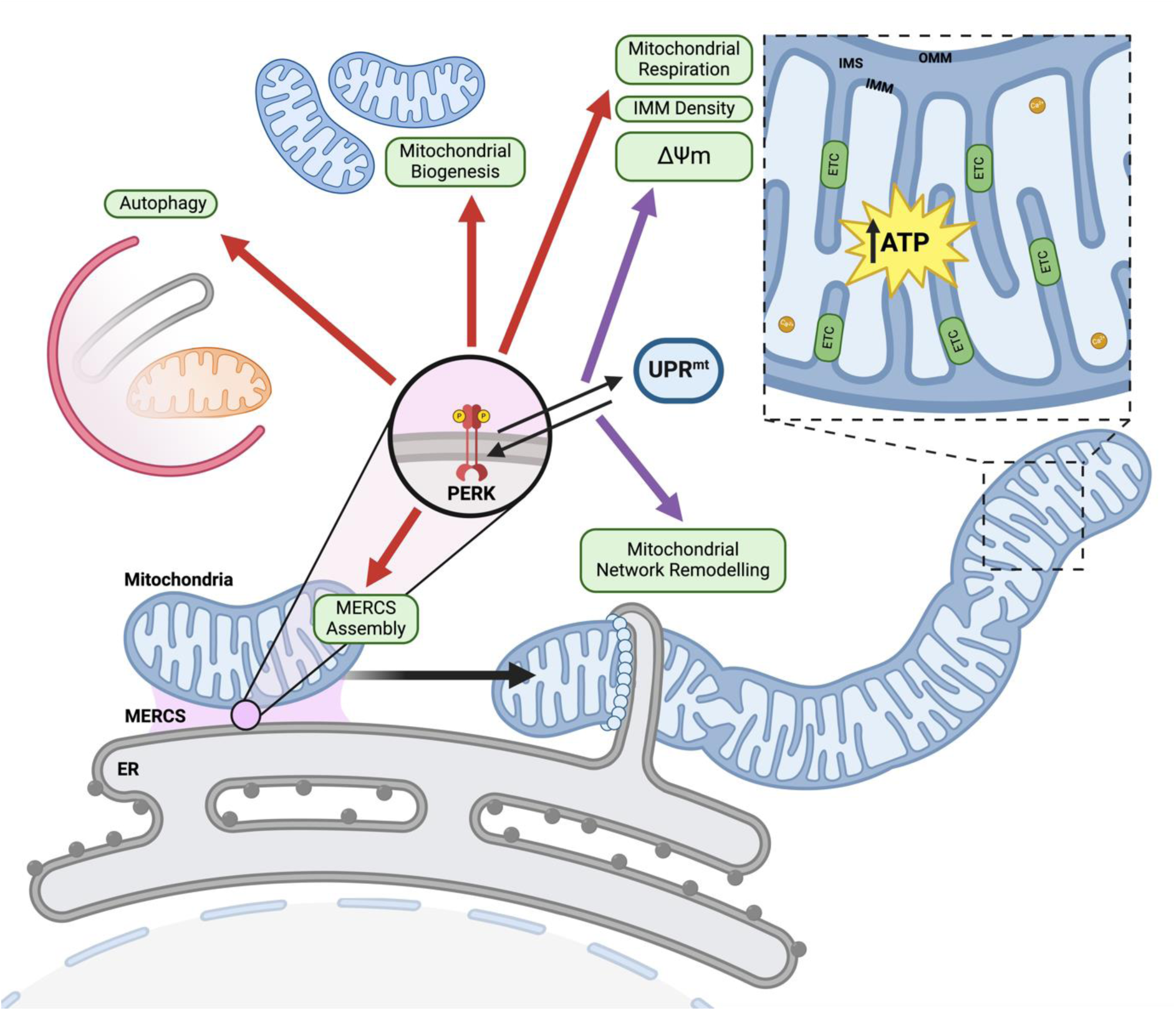

## Introduction

The ER and mitochondria are highly dynamic structures continuously undergoing structural remodelling in response to specific cellular signals. Communication between these organelles can result in physical interactions, called mitochondrial ER contact sites (MERCS), that synergistically affect both organelles ^1,2^. MERCS are dynamic structures that remodel in response to intra and extra cellular signals, affecting the function of both mitochondria and ER ^3–5^. As a result, mitochondrial function is highly sensitive to changes in ER homeostasis and ER stress, which is transmitted to mitochondria through changes in metabolite communication (Ca^2+^ signalling or lipid exchange) or via stress-responsive signalling pathways ^6^. An accumulation of misfolded proteins within the ER can activate the unfolded protein response (UPR^ER^). The activation of the UPR^ER^ can be an adaptive beneficial response, or under conditions of chronic ER stress; it can be maladaptive, resulting in pro-apoptotic signalling events. Disruption of MERCS assembly and communication between these organelles has been reported in an array of age-related diseases including cancer neurodegenerative conditions and sarcopenia ^7^. Targeting or modulating inter organelle communication by modulating MERCS assembly is a potential therapeutic strategy for various pathophysiological conditions.

The UPR^ER^ is a dynamic signalling pathway that is activated following ER stress, mediated by protein kinase RNA-like ER kinase (PERK), inositol-requiring enzyme 1α (IRE1α) and activating transcription factor 6 (ATF6) ^8^. Activation of the PERK arm results in phosphorylation of Serine51 of eukaryotic initiation factor 2α (eIF2α) ^9^, promoting a rapid attenuation of global mRNA translation, reducing the protein load for folding in the ER ^10^. Phosphorylated eIF2α increases selective translation of activating transcription factor 4 (ATF4), inducing the translation of ER stress genes related to the restoration of cellular homeostasis: protein synthesis, amino acid metabolism, redox homeostasis, apoptosis and autophagy ^8^. Accumulation of misfolded proteins, protein aggregation and inefficiency in mitochondrial import, results in the activation of the UPR^mt^ by ATF4, activating transcription factor 5 (ATF5) and C/EBP homologous protein (CHOP) ^11^. Together they promote the transcription of genes that aid in the recovery of normal proteostasis, upregulating chaperonins, chaperones, proteases and antioxidant proteins ^12^. Mitochondria are signalling organelles for a variety of cellular processes including bioenergetics ^13^. The mitochondrial network is dynamic ^14^, the morphology of the mitochondrial network can switch from tubular to hyperfused or fragmented, depending on cellular metabolic conditions and in response to stress insults ^14^. Mitochondrial dynamics maintain the efficient diffusion of metabolites and proteins ^15^, mtDNA levels ^16^, and mitochondria homeostasis ^17^.

PERK is a signal transducer within the UPR^ER^, that can modulate mitochondrial proteostasis and function in response to ER stress and is reported to localise at MERCS ^2,18,19^. PERK activation following ER stress has been demonstrated to be required for acute stress-induced mitochondrial hyperfusion (SIMH) ^18^. Depending on the severity of the cellular stress, ER-mitochondria signalling can directly affect mitochondrial function leading to either pro-survival or pro-apoptotic signals ^20^. An increase in MERCS assembly facilitates Ca^2+^ transfer between the organelles affecting mitochondrial metabolism due to many Ca^2+^-dependent enzymes within the tricarboxylic acid cycle ^21^, increasing mitochondrial ATP generation during adaptive UPR. Prolonged Ca^2+^ entry to mitochondria disrupts mitochondrial dynamics and promotes mitochondrial fragmentation, ultimately leading to the opening of the mPTP, triggering pro-apoptotic signalling ^21^. Mitochondrial function is regulated by fusion, fission and turnover, which determine the organisation of the mitochondrial network ^22^. The ER can coordinate mitochondrial dynamics by establishing contact sites between ER tubules and mitochondria ^23^.

The nematode *Caenorhabditis elegans* is an excellent physiological model for dissecting the mechanisms of adaptive signalling. During ageing, *C. elegans* body wall muscles undergo a decline in mitochondrial dynamics, accumulation of lipids and loss of muscle mass, similar to the ageing process of skeletal muscle from vertebrates ^24^. The UPR^ER^ is highly conserved in nematodes, the three primary signal transducers (PERK, IRE1α and ATF6) and their function resembles their mammalian orthologues ^25^ (Suppl. Fig.1C). The activation of UPR^mt^ in *C. elegans* is regulated by the transcription factor ATFS-1 (ATF5 ortholog in *C. elegans)*. ATFS-1 is imported into the mitochondrial matrix by recognition of its mitochondrial targeting sequence by TOM/TIM, where it is degraded by the matrix-localized protease LON ^11^. Perturbations of mitochondrial homeostasis impair ATFS-1 mitochondrial import and promotes the nuclear translocation of ATFS-1, resulting in the activation of the UPR^mt^ ^26^.

**Figure 1:**
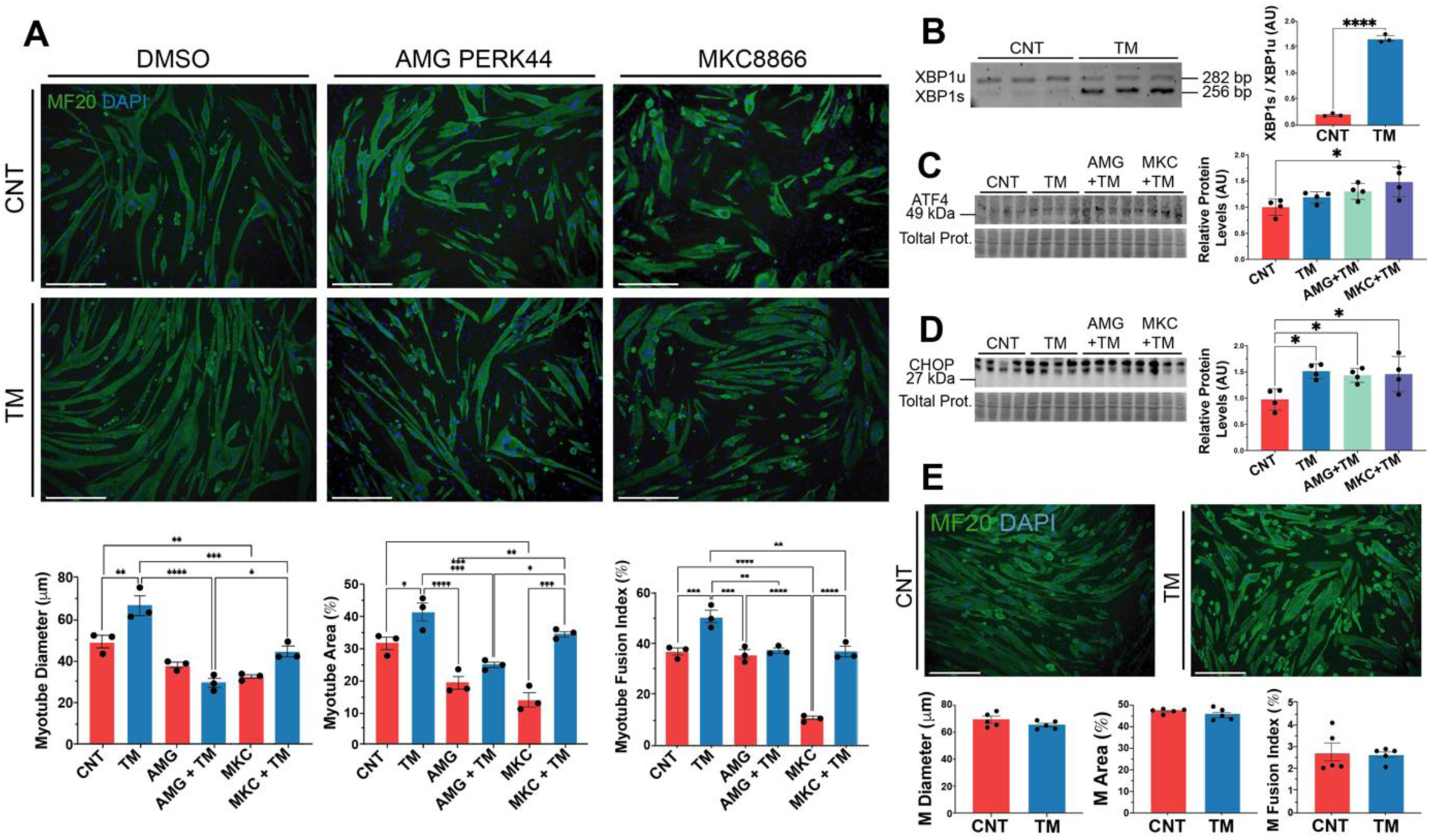
Treatment of C2C12 myoblasts with an acute low concentration of the ER stressor tunicamycin (TM) promotes myogenesis. **A)** MF20 immunostaining following TM treatment of myoblasts resulted in an increase in myotube diameter, area and fusion index. Mean +/- SEM; * p ≤ 0.05, ** p ≤ 0.01, *** p ≤ 0.001 & **** p ≤ 0.0001 One-way ANOVA. **B)** mRNA expression levels of XBP1s/XBP1u in TM-treated myoblasts. Mean +/- SEM; **** p ≤ 0.0001 Student’s t-test. **C & D)** Western blot of ATF4 and CHOP in TM treated myoblasts. Mean +/- SEM; * p ≤ 0.05 & ** p ≤ 0.01 One-way ANOVA. **E)** MF20 immunostaining following TM treatment of myotubes, n=3. Scale 275μm. Mean +/-; Student’s t-test.

This study investigated the role of adaptive UPR^ER^ signalling at an early developmental stage in a cell model of myogenesis and using the whole organism *C. elegans*. Mechanistically adaptive UPR^ER^ signalling resulted in increased MERCS formation, increased mitochondrial dynamics and function that improved myogenesis and healthspan of *C. elegans*. PERK/PEK-1 was identified as the signalling mechanism underlying the adaptive UPR^ER^ response and resulted in downstream activation of ATF5/ATFS-1 required for cellular adaptations. These data provide novel insights into the potential of targeting UPR^ER^ signalling and MERCS assembly. Thereby enhancing the potential to develop therapeutic strategies to mitigate age-related conditions and neurodegenerative diseases, improve overall mitochondrial function and cellular health.

## Results

### PERK promotes myogenesis in myoblasts subjected to physiological levels of ER stress

To investigate the role of adaptive UPR^ER^ during myogenesis, C2C12 myoblasts were exposed to a range of concentrations of the ER stressor tunicamycin (TM) for 8h, followed by differentiation into myotubes (Suppl. Fig.1A). Notably, treatment of myoblasts with 0.2 μg/ml TM significantly enhanced the myogenic potential of myoblasts. However, concentrations above 0.4 μg/ml were detrimental and promoted cell death (Suppl. Fig.2A). To determine if the response was an ER stress response driving activation of UPR or specific to TM, the SERCA inhibitor thapsigargin (TG) was also used. A low concentration of TG (10nM) treatment for 8h also promoted an increase in myogenic parameters (Suppl. Fig.2B). Inhibition of PERK with AMG PERK44 (AMG) resulted in no difference in myogenesis parameters compared to controls (Fig.1A). Furthermore, the combined AMG+TM treatment group (AMG+TM) exhibited no change when compared to the AMG group (Fig.1A). However, inhibition of IRE1α with MKC8866 (MKC) led to a significant decrease in all the parameters compared to controls (Fig.1A). Nevertheless, the MKC+TM group exhibited an increase in myogenic parameters compared to MKC alone (Fig.1A). Interestingly, TM-treated myoblasts exhibited a significant decrease in cell viability, with an even greater decrease observed in the AMG+TM group (Suppl. Fig.2C).

**Figure 2:**
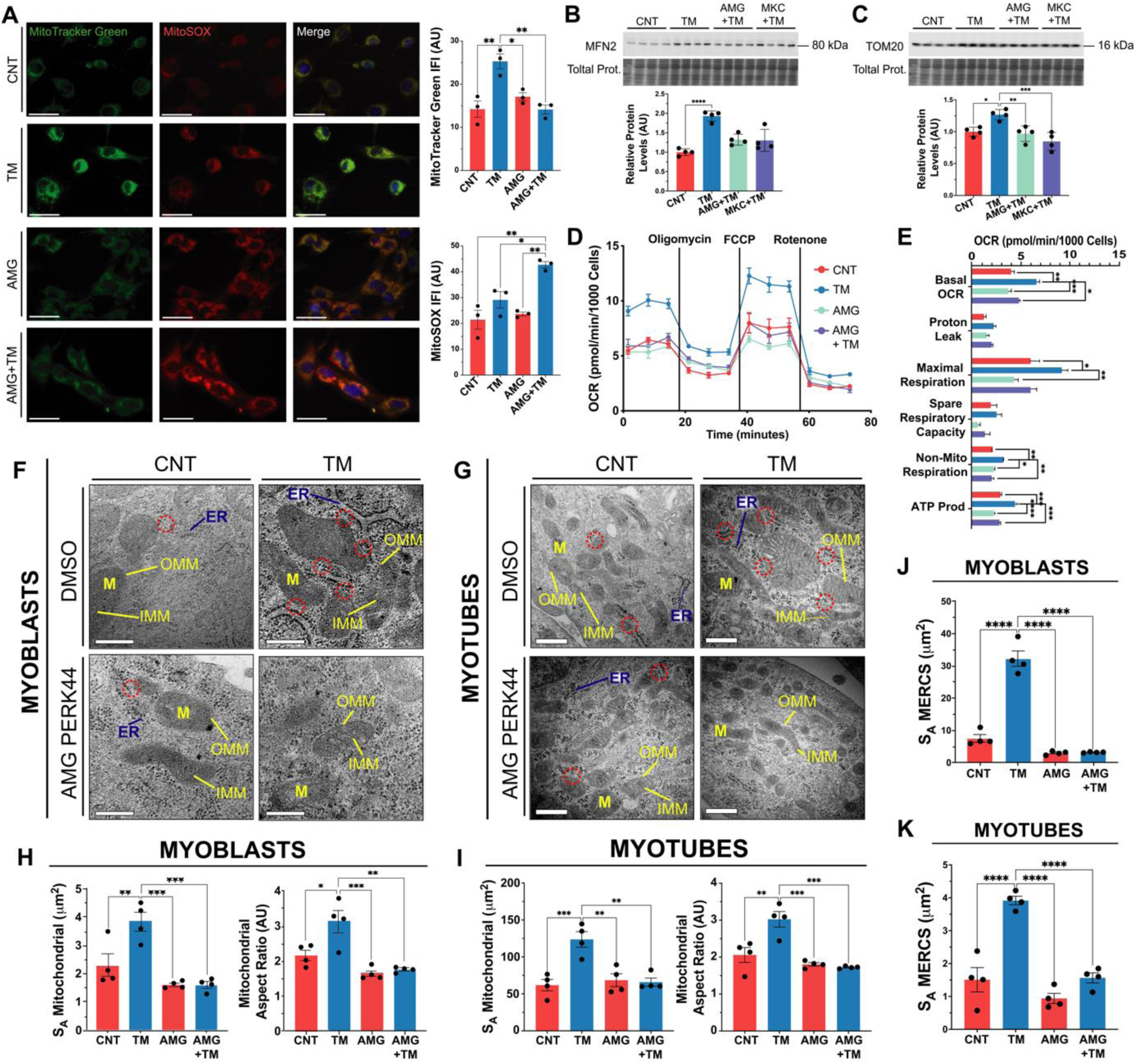
Adaptive UPR^ER^ signalling increases mitochondrial capacity and MERCS assembly depends on the PERK arm of the UPR^ER^. **A)** TM treatment increased mitochondrial content (MitoTracker Green) but did not affect MitoSOX staining; n=3. Scale 40μm. Mean +/- SEM; * p ≤ 0.05 & ** p ≤ 0.01 One-way ANOVA. **B, C)** Western blot of MFN2 and TOM20 in TM treated myoblasts. Mean +/- SEM; * p ≤ 0.05 & ** p ≤ 0.01 One-way ANOVA. **D, E)** TM treatment of myoblasts increased basal and maximal respiration rate, ATP production and non-mitochondrial respiration. Mean +/- SEM; * p ≤ 0.05, ** p ≤ 0.01, *** p ≤ 0.001 & **** p ≤ 0.0001 One-way ANOVA. **F, H & J)** Representative TEM images of sections of myoblasts. Mitochondrial surface area, aspect ratio and MERCS surface area increased with the TM treatment. PERK inhibition resulted in no differences compared to the control.; n=4. Scale 200nm. Mean +/- SEM; * p ≤ 0.05, ** p ≤ 0.01, *** p ≤ 0.001 & **** p ≤ 0.0001 One-way ANOVA. **G, I & K)** Representative TEM images of sections from myotubes following TM treatment of myoblasts demonstrating the increase in mitochondrial surface area, aspect ratio and MERCS surface area was maintained in myotubes following TM treatment.; n=4. Scale 200nm. Mean +/- SEM; * p ≤ 0.05, ** p ≤ 0.01, *** p ≤ 0.001 & **** p ≤ 0.0001 One-way ANOVA.

PCR analysis confirmed activation of the IRE1α arm of the UPR, with a shift toward the spliced, active form of XBP1 (XBP1s) following TM treatment (Fig.1B). Western blot analysis of UPR markers revealed no significant changes in GRP78 levels, though there was an increasing trend in eIF2α phosphorylation (Suppl. Fig.2D). There was an increase in the levels of ATF4 in MKC+TM-treated cells, while CHOP increased following TM treatment and also elevated in both AMG+TM and MKC+TM groups (Fig.1C&D). Similar to inhibition of PERK with AMG44, knock-down of PERK using siRNA (siPERK) in myoblasts decreased myotube diameter following differentiation. Knock-down of PERK or CHOP combined with the TM treatment, exhibited a reduction in myotube area and fusion index compared to siRNA CNT (Suppl. Fig.2E). Additionally, TM treatment of differentiated, untreated myotubes did not result in any significant differences in the myotube parameters (Fig.1E). Collectively, these findings demonstrate that inducing a mild ER stress in myoblasts can improve myogenesis and identified an essential role of PERK in adaptive UPR signalling.

### PERK promotes MERCS assembly and mitochondrial adaptations during adaptive UPR

The effects of TM treatment (0.2μg/ml for 8h) and/or PERK inhibition on mitochondrial content and function were evaluated in myoblasts (Fig. 1A). MitoTracker Green staining was used to assess mitochondrial content and increased with TM treatment. No differences were observed between the AMG group and the controls (Fig.2A). Mitochondrial ROS was assessed using MitoSOX staining and there was no change in MitoSOX staining in the TM and AMG groups. However, an increase in fluorescence was observed in the AMG+TM group (Fig.2A). Additionally, MitoTracker Green and TMRE staining was used to assess mitochondrial membrane potential (ΔΨm), there was a significant increase in both MitoTracker Green and TMRE staining following TM treatment. The ratio of the intensities indicated that TM treatment promoted mitochondrial biogenesis and enhanced ΔΨm (Suppl. Fig.2F). Both the mitochondrial fusion protein MFN2, which plays a role in mitochondrial-ER tethering ^27,28^, and the mitochondrial content indicator TOM20 increased with TM treatment (Fig.2B&C). The oxygen consumption rate of C2C12 myoblasts demonstrated enhanced basal and maximal respiration, non-mitochondrial respiration, and ATP production in TM-treated cells compared to controls, AMG, and AMG+TM groups (Fig.2D&E). Furthermore, transmission electron microscopy (TEM) organelle ultrastructure analysis of myoblasts demonstrated an increase in the membrane surface area of mitochondria (S_A_), mitochondrial elongation (aspect ratio) following TM treatment; these adaptations were blocked by PERK inhibition (Fig.2F&H). Notably, these mitochondrial adaptations were preserved following differentiation into myotubes (Fig.2G&I). Additional parameters, including mitochondrial volume fraction, cristae membrane density (S_A_ IMM/OMM) and ER surface area, increased in both myoblasts and subsequent myotubes following TM treatment (Suppl. Fig.2G). The evaluation of MERCS surface area demonstrated an expansion of mitochondrial and ER membranes that are in close proximity following TM treatment. The formation of these expanded subdomains was blocked by inhibition of PERK (Fig.2F&J). The increase in MERCS surface area was sustained after differentiation into myotubes (Fig. 2G&K). Together, the data demonstrate that PERK is required during adaptive UPR^ER^ to form MERCS and increase mitochondrial content and function in myoblasts, which were maintained following differentiation into myotubes.

### PERK is required for adaptive UPR^ER^ activation of autophagy and mitophagy

To determine the mechanisms underlying the adaptations promoted by the UPR^ER^ in myoblasts, proteomic changes induced by an 8h treatment with 0.2 μg/ml TM using label-free proteomics were determined. Global label-free proteomics analysis quantified 2513 proteins, 71 proteins were significantly upregulated and 27 downregulated following TM treatment (Fig.3A). Enrichment analysis of the 71 significantly upregulated proteins identified several pathways and proteins involved in *Mitophagy* and *mitochondrial ADP/ATP transport* (ADRO, TPP1, TXTP, MTDC, SYHM, RM12, KAPCA, ADT1 and ADT2), indicating active remodelling of the mitochondrial network and mitochondrial respiration. Upregulated proteins related to *Cell cycle and differentiation* (PTBP3, RCC1, CCNK, MD1L1 and MYH10), suggest enhanced myogenesis. Proteins linked to the *Cytoskeleton* (CKAP5 and CKAP4), indicate an increased regulation of organelle contact and communication (Fig.3B). No significant pathway enrichment was identified among the 27 downregulated proteins, which spanned diverse biological processes, including *mitochondrial RNA processing* and *protein translation* (4EBP1, MRT4, RPR1B and EI2BA) (Fig.3A).

**Figure 3:**
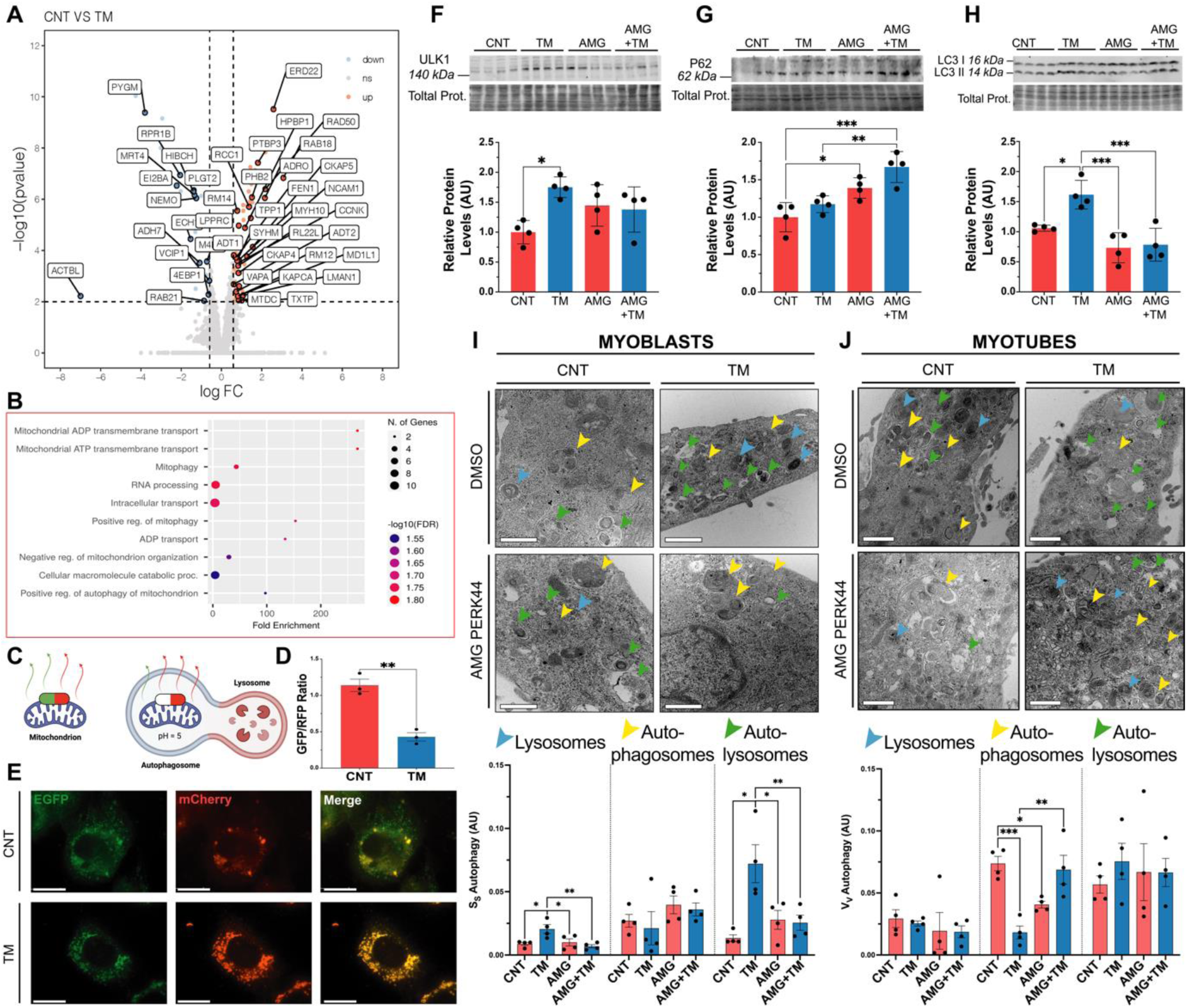
Adaptive UPR^ER^ signalling in myoblasts promotes autophagy machinery and mitochondrial turnover. **A)** Volcano plot of LFQ proteomic data of protein abundance following TM treatment (FDR=1.58%). **B)** GOBP ShinyGO enrichment of significantly upregulated proteins. **C)** Graphical diagram of the pH-sensitive construct for visualisation of mitochondrial turnover. **D & E)** Increased mitophagy in TM treated myoblasts, represented by a decrease in the GFP/ RFP ratio.; n=3. Mean +/- SEM; ** p ≤ 0.01 Student’s t-test. Scale 25μm. **F, G & H)** Western blot of protein extracts for ULK1, p62 and LC3 II/I. Error bars SEM; * p ≤ 0.05, ** p ≤ 0.01 & *** p ≤ 0.001 One-way ANOVA. **I)** Representative TEM images of sections of TM treated myoblasts and quantification of the lysosome, autophagosomes and autolysosome fractions. n=4. Scale 400nm. Mean +/- SEM; * p ≤ 0.05 & ** p ≤ 0.01 One-way ANOVA. **J)** Representative TEM images of sections of myotubes treated with TM at myoblast stage and quantification of lysosome, autophagosome and autolysosome fractions. n=4. Scale 400nm. Mean +/- SEM; * p ≤ 0.05, ** p ≤ 0.01 & *** p ≤ 0.001 One-way ANOVA.

To further explore mitochondrial turnover, live-cell imaging was performed using the mitophagy reporter plasmid [*Cox8-EGFP-mCherry*], transfected into myoblasts before TM treatment. The reporter contains a pH-sensitive GFP and a pH-stable RFP to track mitochondrial turnover; GFP fluorescence is quenched in acidic lysosomes, while RFP remains stable, allowing an estimation of mitophagy (Fig.3C). TM treatment significantly reduced the GFP/RFP ratio compared to the control group, indicating an induction of mitophagy (Fig.3D &E). Western blot analysis of autophagy-related proteins revealed that ULK1 levels were elevated in the TM group compared to the CNT, AMG, and AMG+TM groups (Fig.3F). p62 levels increased in the AMG and AMG+TM groups compared to the CNT and TM groups, indicating reduced autophagic flux (Fig.3G). Finally, increased expression of ATG5 (Suppl. Fig.2H) and LC3-II/I (Fig.3H) in the TM group suggested enhanced autophagosome formation. Ultrastructure analysis of the autophagic bodies by TEM, evaluated changes in the volume fraction of lysosomes, autophagosomes and autolysosomes from myoblasts and myotubes. Myoblasts exhibited an apparent increase in the volume fraction of lysosomes and autolysosomes in the TM group compared to CNT, AMG and AMG+TM groups (Fig.3G). In contrast, myotube analysis indicated no change in lysosome or autolysosomes volume fractions between groups, although there was a decrease in autophagosome volume in the TM group compared to the CNT and AMG+TM groups (Fig.3H). Changes in [Ca^2+^]_i_ showed a slight, but not significant, increase in the maximal increase of store-operated calcium entry (SOCE) from the TM-treated myoblasts compared to controls (Suppl. Fig.2I). Additionally, ER volume was assessed using an ER-targeting plasmid [*ER (KDEL)-mNeonGreen*] to estimate the ER membrane area relative to cellular surface area, revealing a significant expansion in ER membrane area following TM treatment (Suppl. Fig.2J). Overall, proteomic and ultrastructural analyses revealed that TM treatment induces mitophagy and enhances autophagic flux during myogenesis. These adaptations depend on PERK activity and potentially play a crucial role in regulating organelle turnover.

### Early-life induction of the UPR^ER^ extends lifespan and preserves healthspan in C. elegans

*C. elegans* was used as a physiological model to assess the role of UPR^ER^ signalling in a whole organism. To replicate the *in vitro* myogenesis model, worms were treated at an early developmental stage. Following bleaching, embryos were harvested and exposed to 1.25 μg/ml TM in S Medium for 24h and larvae plated and allowed to develop normally (Suppl. Fig.1B). There was an increase in the longevity of TM-treated worms compared to controls (Fig.4A). Reproductive potential was significantly reduced in the TM group, particularly in the later stages of the reproductive period (Fig.4B). Importantly, this decrease in progeny was not associated with increased embryo lethality Suppl. Fig.3A). Interestingly, TM-treated worms exhibited increased body length compared to controls (Fig.4C). The activation of both the UPR^ER^ and the UPR^mt^ was confirmed using transcriptional GFP reporter strains for *hsp-4* (orthologue of mammalian HSPA5/Grp78) and *hsp-6p* (ortholog of the human mtHSP70). TM treatment resulted in greater UPR^ER^ and UPR^mt^ activation capacity compared to controls (Fig.4D&E). Analysis of the IRE-1 branch of the UPR revealed increased splicing of XBP-1 in TM-treated worms (Fig.4F). The survival of the worms against a range of different toxicants was assessed. The resistance of the TM treated group against sodium arsenite, the redox cycler Paraquat and the organic peroxide tBuOOH was higher than the control group at adult day1 (Fig.4G-I). CeleST (*C. elegans* Swim Test) was performed to assess physical fitness and mobility during ageing ^29^. The fitness levels were evaluated on days 1, 5, 10 and 15 of adulthood. Fitness markers including wave initiation rate, travel speed and activity index decreased with age in both groups but remained significantly higher in TM-treated worms compared to controls at all time points (Fig.4J, Suppl. Fig.3E&F). Conversely, frailty markers including stretch, average body curvature and curling increased with age in both groups, but these were consistently lower in the TM group, indicating better maintenance of physical health in the TM-treated groups (Fig.4K, Suppl. Fig.3G&H). These results suggest that early-life UPR^ER^ activation extends lifespan, increases stress resistance and also promotes healthspan by preserving physiological fitness and reducing frailty during ageing.

**Figure 4:**
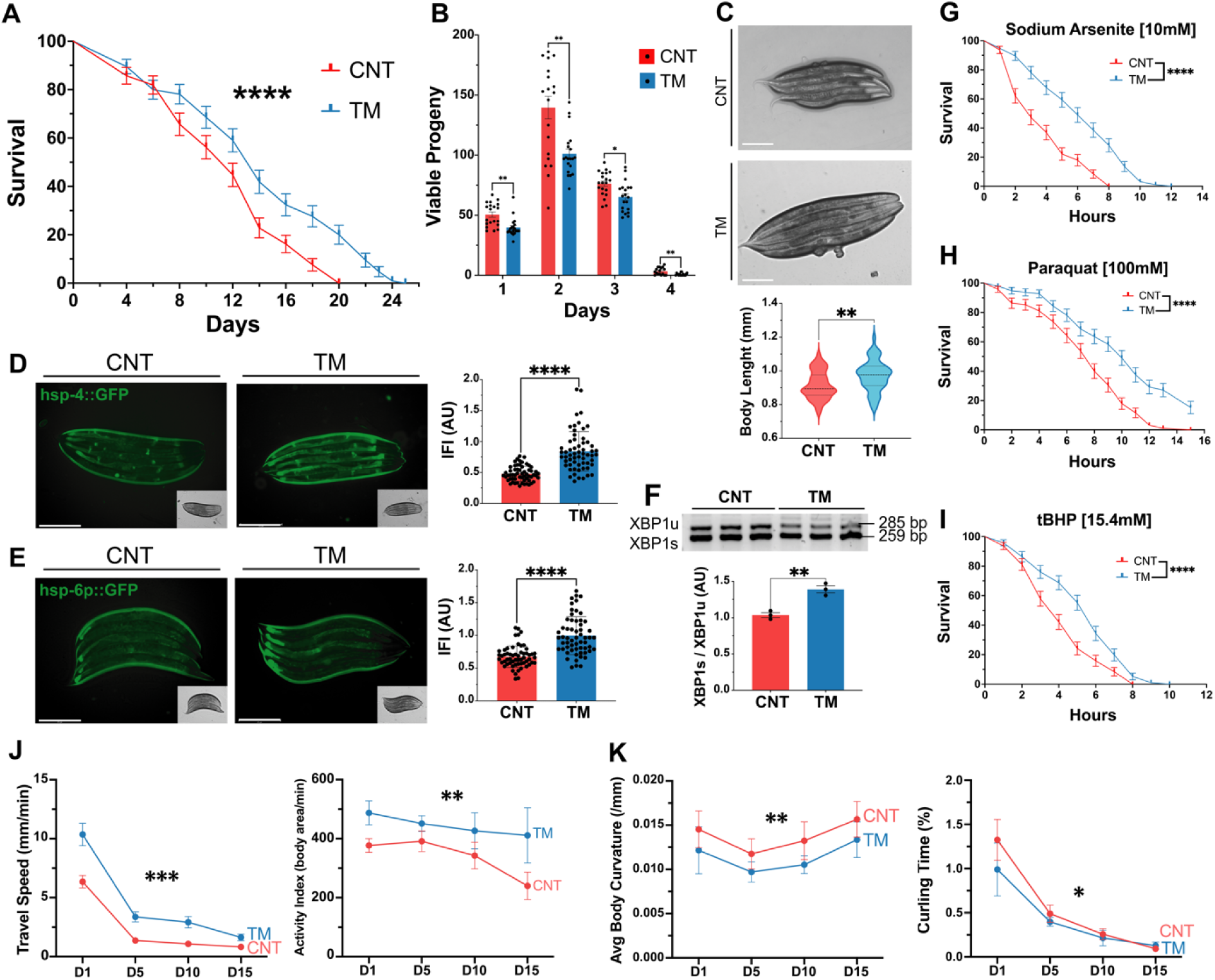
Activation of UPR^ER^ in *C. elegans* at an early developmental stage increased longevity, stress resistance and fitness. **A)** Lifespan assay of TM treated N2 strain. Kaplan–Meier survival plots of two independent experiments initiated with 100 animals/group. Mean +/- SEM; **** p ≤ 0.0001 Log-rank (Mantel-Cox) test. **B)** TM treatment decreased progeny in N2 wild-type *C. elegans*. Data from 20 animals/group. Mean +/- SEM; * p ≤ 0.05 & ** p ≤ 0.01 Student’s t-test. **C)** Body length of N2 wild-type TM-treated worms. Data from 20 animals/assay. Mean +/- SEM; ** p ≤ 0.01 Student’s t-test. **D, E)** Activation of UPR assessed by *hsp-4*::GFP (UPR^ER^)and *hsp-6p*::GFP (UPR^mt^) following TM treatment. Scale 275μm. Mean +/- SEM of at least 20 animals/assay. **** p ≤ 0.0001 Student’s t-test. **F)** RT-PCR analysis of expression ratio between XBP1s:XBP1u transcripts. n=3, Error bars SEM; ** p ≤ 0.01 Student’s t-test. **G, H & I)** Survival of wild type in Sodium Arsenite, tBuOOH and Paraquat. Kaplan–Meier survival plots of two independent experiments initiated with 45 animals/ group. Mean +/- SEM; **** p ≤ 0.0001 by Log-rank (Mantel-Cox) test compared with wild-type control. **J)** CeleST physical fitness parameters at days 1, 5, 10, & 15. Data mean of at least 30 animals/assay. **K)** CeleST physical frailty parameters at days 1, 5, 10, & 15 individual values controls and TM-treated worms during ageing. Data mean of at least 30 animals/assay.

### Early-life induction of the UPR promotes lifespan extension in C. elegans through modulation of lipid metabolism and lysosome activation

As TM treatment of myoblasts and myotubes resulted in very different effects on myotubes, we assessed whether the adaptations promoted by the UPR^ER^ during early nematode development were similar to TM treatment of adult worms. TM treatment (1.25 and 5μg/ml for 24h) was performed during L4 to adult Day1 developmental stage of *C. elegans*. Both TM treatments (1.25 and 5μg/ml) at L4 stage resulted in a significant reduction in lifespan (Fig.5A), along with decreased resistance to paraquat and sodium arsenite (Fig.5B&C). Interestingly, no change in resistance to sodium arsenite was observed in worms treated with the lower TM dose (1.25μg/ml) (Fig.5B). This highlights that the beneficial effects of TM treatment occurred only at an early developmental stage and treatment of adult worms resulted in negative effects similar to the myogenesis model.

**Figure 5:**
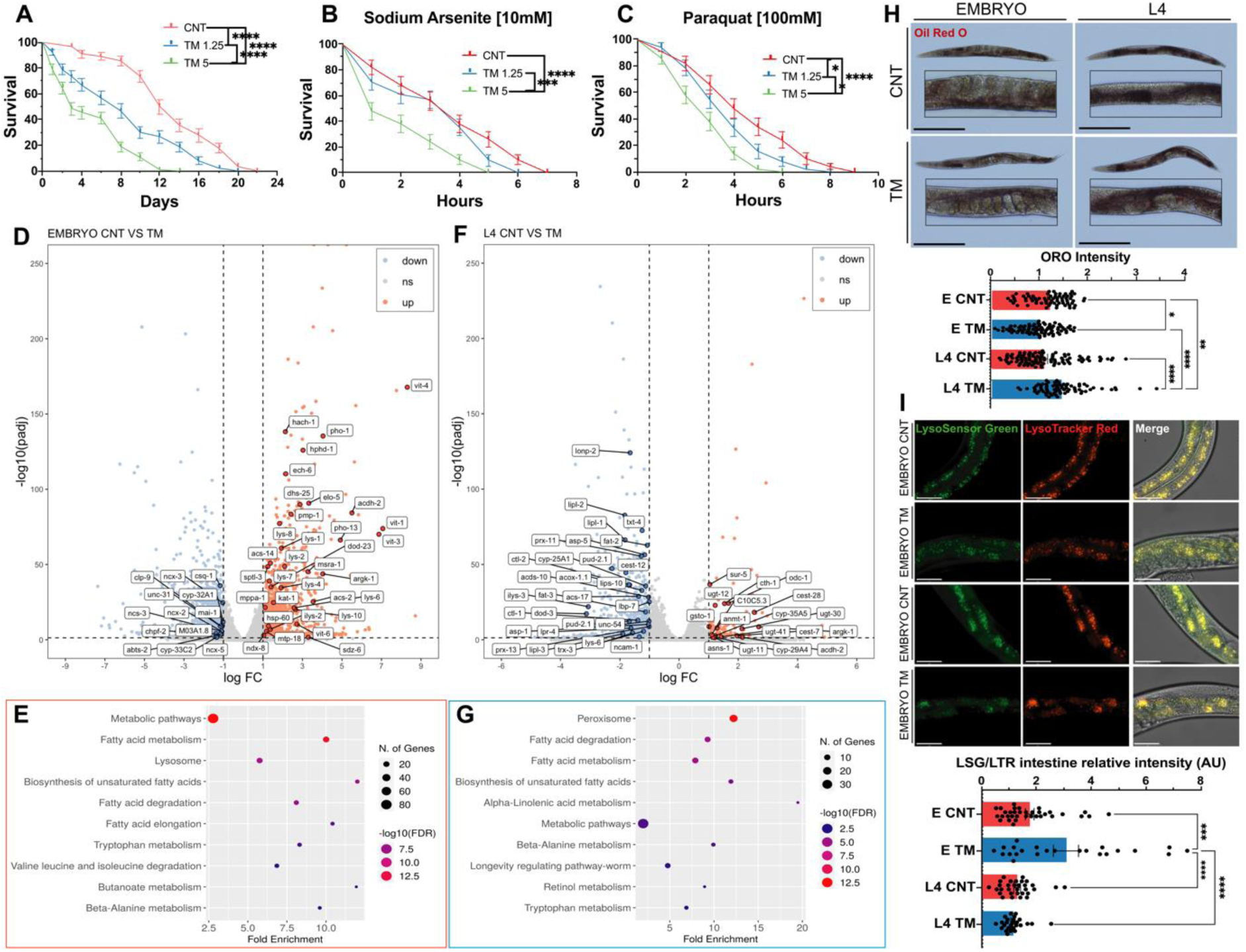
UPR^ER^ activation in *C. elegans* embryos or at L4 stage has opposite effects on lipid metabolism. **A)** Lifespan analysis of N2 strain treated with TM at L4 stage. Kaplan–Meier survival plots of two independent experiments initiated with 100 animals. Mean +/- SEM; **** p ≤ 0.0001 Log-rank (Mantel-Cox). **B & C)** Decreased survival of N2 worms treated with TM at L4 stage in 10 mM sodium arsenite and 100 mM Paraquat. Kaplan–Meier survival plots of two independent experiments initiated with 50 animals/group. Mean +/- SEM; * p ≤ 0.05, *** p ≤ 0.001 & **** p ≤ 0.0001 Log-rank (Mantel-Cox). **D & F)** Volcano plot of differentially expressed genes (DEGs) following TM treatment of embryos (D) or adults (F), red significantly upregulated genes and blue significantly downregulated genes. 4 biological replicates per condition. **E & G)** Dot plot of Shiny GO KEGG analysis of DEGs. Upregulated pathways are in red and downregulated pathways are in blue. **H)** Oil Red O stained N2 worms following TM treatment at embryo (E) or L4 stage. 3 independent experiments with at least 20 animals per strain; Scale 275μm. Mean +/- SEM; * p ≤ 0.05, ** p ≤ 0.01 & **** p ≤ 0.0001 by ANOVA. **I)** Intestine of wild-type worms incubated with LysoSensor Green (LSG) indicator of lysosomal pH and LysoTracker Red (LSR) indicator of lysosome content, 3 independent experiments with 10 animals per condition; Scale 50μm. Mean +/- SEM; *** p ≤ 0.001 & **** p ≤ 0.0001 by ANOVA.

RNA sequencing was performed to determine the molecular mechanisms underlying the different responses dependent on the developmental stage. A transcriptome analysis was conducted on Day1 adults that were treated with TM 1.25μg/ml at embryonic stage (EMBRYO group) (Fig.5D-F) or treatment with TM at L4 stage (L4 group) (Fig.5G-I). Differentially expressed genes (DEGs) were analysed (Log_2_-fold change > 1 and adjusted p-value<0.05) from EMBRYO (FDR=2.38%) and L4 separately (FDR=1.1%). DEGs from EMBRYO-treated groups reported 1249 upregulated and 2042 downregulated genes. Functional enrichment analysis of the upregulated genes identified pathways related to lipid metabolism and lysosome activation (Fig.5E). Both pathways have previously been described as key intermediates in the UPR regulation of ageing ^30^. Gene set enrichment analysis (GSEA) indicated enhanced expression of genes associated with autophagy (lysosomal pathways) and mitochondrial activity (oxidative phosphorylation) (Suppl. Fig.3I). Lipid biosynthesis emerged as the most prominent upregulated pathway, and there was a marked increase in most intermediates of this pathway (Suppl. Fig.3K). In contrast, the L4 group resulted in 146 upregulated and 811 downregulated genes (Fig. 5G). Functional enrichment of the downregulated genes highlighted pathways involved in lipid metabolism and *C. elegans* longevity (Fig.5H). GSEA revealed decreased expression of genes related to autophagy (lysosomes) and mitophagy in L4-treated worms (Suppl. Fig.3J). Pathway enrichment analysis identified a reduction in the expression of intermediates in the lipid biosynthesis pathway in the L4 group (Suppl. Fig.3L). Finally, comparing the functional enrichment of downregulated genes in the EMBRYO group with the upregulated genes in the L4 group demonstrated shared pathways related to metabolism (Suppl. Fig. 3B & C). These findings suggest that UPR^ER^ activation during early development promotes beneficial adaptations that contribute to lifespan extension, whereas induction later in life leads to a decline in similar protective mechanisms, particularly affecting lipid metabolism and autophagy.

RNA sequencing highlighted the effects of TM treatment on lipid metabolism and the lysosome. Oil Red O staining, which stains neutral triglycerides and lipids, was performed in the different groups ^31^. Interestingly, TM treatment decreased lipid staining in the EMBRYO group compared to the controls (Fig.5J). However, in the L4 group, TM treatment increased lipid staining compared to the control group (Fig.5J), supporting the transcriptomic data of increased expression of lipid-related genes in embryo-treated worms but a decrease in L4 treated worms. The lysosomal content and acidity were estimated using the LysoSensor green/LysoTracker red (LSG/LSR) ratio in the intestine ^32^. Results demonstrated decreased pH in the lysosomes of N2 worms subjected to the TM treatment at the embryonic stage. No changes in lysosomal pH were reported in the animals treated at the L4 stage (Fig5K). An upregulation of lysosomal genes and increased intestinal lysosomal acidity in response to UPR^ER^ activation has previously been demonstrated to improve proteostasis and longevity in *C. elegans* ^33^. Overall, the effects of TM treatment were dependent on the developmental stage and resulted in the opposite expression of genes involved in lipid metabolism and subsequent effects on longevity and oxidative stress resistance. The results support the findings in the myogenesis model and highlight increased plasticity for adaptations following UPR^ER^ induction at early developmental stages.

### Early-life UPR induction stimulates MERCS formation, mitochondrial capacity and mitophagy in C. elegans

In order to determine if there was a similar adaptive response in nematodes to the myogenesis model following TM treatment, the effects on mitochondrial content was assessed using a transcriptional P*cox-4*::GFP reporter ^34^. The fluorescence intensity increased with TM treatment (Fig.6A). MitoTracker Red staining was used to estimate mitochondrial membrane potential and there was a significant increase in the fluorescence intensity following treatment with TM (Fig.6B). In addition, mitochondrial superoxide production was evaluated by MitoSOX staining, demonstrated a reduction in the TM group (Suppl. Fig.3D). To further examine mitochondrial function, oxygen consumption rate (OCR) was performed using adult day1 *C. elegans* following treatment with TM at the embryo stage. TM-treated worms showed a slight, non-significant increase in both basal and maximal OCR, and there was a significant increase in non-mitochondrial respiration (Fig.6C). TEM was used to analyse the mitochondrial ultrastructure of body wall muscle cells from *C. elegans* (Suppl. Fig.1D). Stereological analysis revealed increased mitochondrial surface area in the TM group (Fig.6D). Moreover, mitochondria from the TM treatment group, exhibited an increased aspect ratio, indicating more elongated, fused mitochondria. The surface area ratio between the inner (IMM) and outer mitochondrial membranes (OMM) was also significantly increased, reflecting a denser IMM structure (Fig.6D), which may contribute to the observed improvements in mitochondrial function.

**Figure 6:**
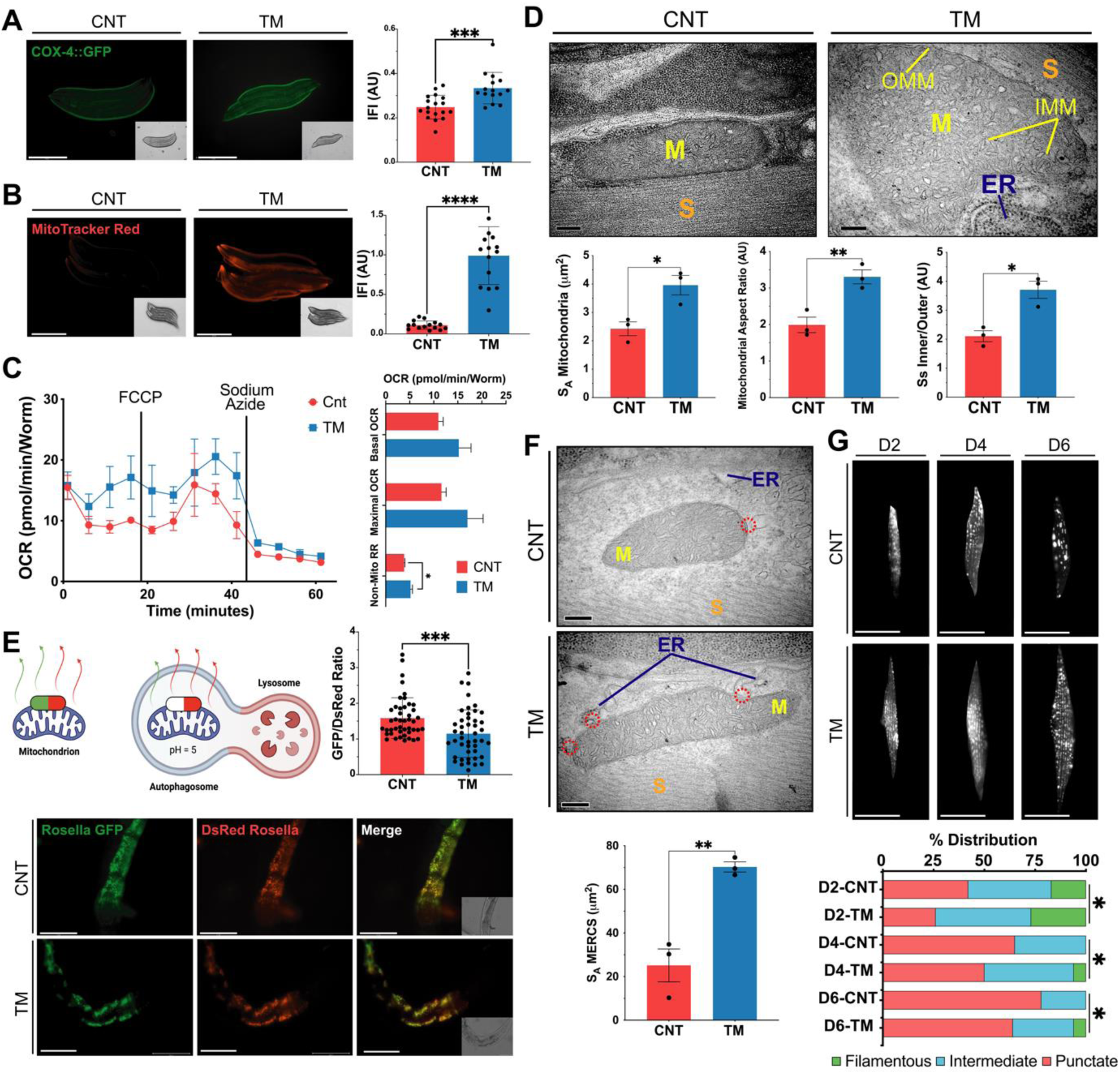
Early-life exposure of *C. elegans* to ER stress improves mitochondrial capacity, activates mitophagy and promotes MERCS assembly. **A)** Mitochondrial content assessed using transgenic worms *pcox-4*::*gfp* following TM treatment. 3 independent experiments with 15 animals/group. Scale 275μm. Mean +/- SEM; *** p ≤ 0.001. Student’s t-test. **B)** MitoTracker red staining of N2 strain following TM treatment. 3 independent experiments were initiated with 15 animals/group. Scale 275μm. Mean +/- SEM; **** p ≤ 0.0001. Student’s t-test. **C)** Oxygen consumption of adult N2 worms following treatment with TM. Data from 3 independent experiments with 8-12 animals per well. Mean +/- SEM; * p<0.05 Student’s t-test. **D)** Representative TEM images of sections from *C. elegans* body wall muscle cells. Mitochondrial surface area, aspect ratio and IMM/OMM surface area ratio were calculated. Scale 200nm. Mean +/- SEM; * p ≤ 0.05 & ** p ≤ 0.01 Student’s t-test. **E)** Representative images of *pmyo-3::TOMM-20::Rosella* reporter strain following TM treatment from 3 independent experiments initiated with 15 animals/group.; ** p<0.01 Student’s t-test. Scale 75μm. **F)** Representative TEM images of sections from *C. elegans* body wall muscle cells. Representative images of 3 independent experiments initiated with 10 animals/group. Scale 200 nm. Mean +/- SEM; ** p ≤ 0.01 Student’s t-test. **G)** Representative images of p*myo-3::mitogfp* reporter strain for muscle mitochondrial morphology, classified as either filamentous, intermediate or punctate. 3 independent experiments were initiated with 15 animals/ condition. Scale 50μm. n=3 * p<0.05 Chi-square.

Mitochondrial turnover was assessed using the *pmyo-3::TOMM-20::Rosella* reporter strain, containing a pH-sensitive Rosella biosensor composed of a pH-sensitive GFP and a pH-stable RFP to track mitochondrial turnover in body wall muscle ^35^. TM-treated worms exhibited a significantly lower GFP/DsRed fluorescence ratio, indicating increased mitophagy and enhanced mitochondrial turnover during adulthood (Fig.6E). As MERCS are essential for mitochondrial fission, TEM analysis was performed to quantify the surface area of mitochondria in close contact with the ER (< 50 nm). TM-treated worms resulted in a marked increase in MERCS surface area compared to controls (Fig.6F). Finally, we evaluated mitochondrial morphology during ageing using a genetic p*myo-3::mitoGFP* reporter, for visualisation of the mitochondrial network from body wall muscle. Images were taken on days 2, 4 and 6 of adulthood and classified as punctate, intermediate or filamentous, depending on the mitochondrial network morphology ^36^. Ageing resulted in a progressive shift towards more punctate (fragmented) networks (Fig.6G). In contrast, TM-treated worms demonstrated a delayed onset of mitochondrial fragmentation, maintaining more filamentous and intermediate networks at all time points assessed (Fig.6G). Together, the data demonstrates that, like myoblasts, early-life TM treatment helps preserve mitochondrial network integrity and morphology during ageing, potentially through the formation of MERCS and regulation of mitochondrial turnover.

### PEK-1 (UPR^ER^) and ATFS-1 (UPR^mt^) crosstalk regulates the adaptations promoted by the UPR in C. elegans

To elucidate the mechanisms underlying UPR-mediated mitochondrial and physiological adaptations, we used mutant strains for the UPR^ER^ and UPR^mt^ signalling arms to evaluate their response to early-life TM treatment. Both the *ire-1(v33)* and *xbp-1(tm2482)* mutant strains were viable; however, TM treatment resulted in developmental arrest at the larval stage (Suppl Fig.3A). This phenotype has previously been reported in *C. elegans*, generation of double mutants of IRE-1 or XBP-1 with ATF-6 or PEK-1 promoted developmental arrest at larvae stage L2 ^37^. This phenotype highlights the critical role of the IRE-1 arm in the resistance to ER stress. Treatment of the *atf-6 (ok551)* mutant strain with TM resulted in increased longevity and stress resistance to sodium arsenite but not paraquat (Suppl. Fig.4B). The TM-treated group had increased MitoTracker Red fluorescence intensity (Suppl. Fig.4B). Following TM treatment, *atf-6* mutants exhibited a reduction in size at day1 of adulthood, implying delayed development (Suppl. Fig.4B). As the increased longevity and survival following TM treatment were also observed in N2 strain (Fig.4A, G&H), it would suggest they are independent of ATF-6 signalling. Following TM treatment of the PEK-1 loss-of-function mutant (*pek-1* (ok275)), there was a decrease in lifespan (Fig.7A), resistance to paraquat was reduced but resistance to sodium arsenite was unaltered (Suppl. Fig.4C). There was a decrease in MitoTracker Red fluorescence intensity in the TM-treated *pek-1* mutants (Fig.7B). The *pek-1* mutant strain also had a reduction in size following TM treatment (Suppl. Fig.4C). TM treatment of *pek-1* mutant strain produced an opposite phenotype to that observed in the N2 strain, highlighting its importance for this adaptive response. TM treatment of ATFS-1 mutant (*atfs-1* (tm4525)) also exhibited a decrease in lifespan (Fig.7C); however, their resistance to oxidative stressors were unaltered (Suppl. Fig.4D). The ΔΨm of the TM-treated *atfs-1* mutant strain decreased following TM treatment (Fig.7D). Interestingly, TM-treated *atfs-1* mutant exhibited no differences in size compared to the control group (Suppl. Fig.4D). The results highlight that TM treatment of the *atfs-1* mutant strain had similar results to the *pek-1* mutant and an opposite phenotype to N2 WT strain, highlighting its importance for this adaptive response.

**Figure 7:**
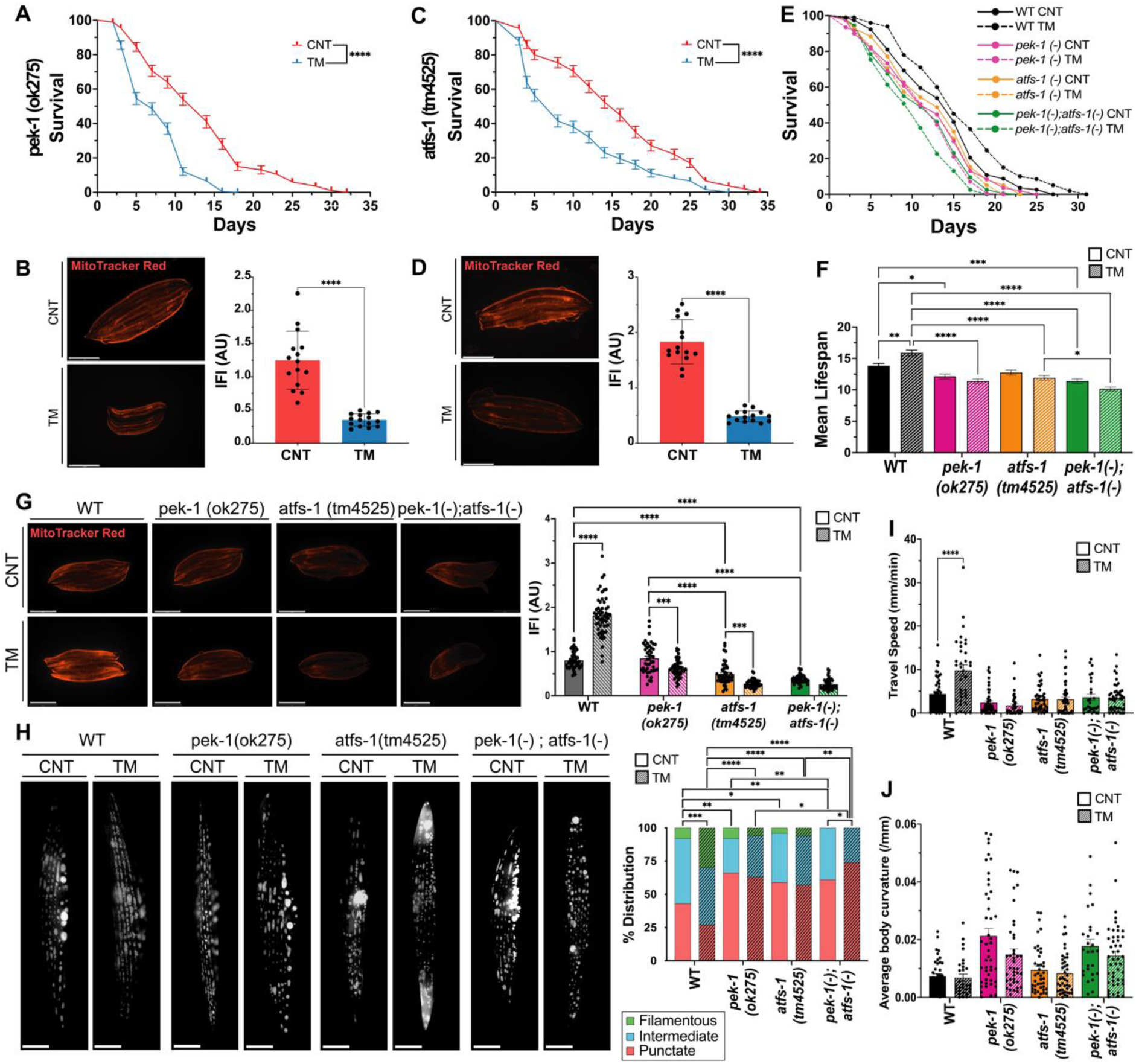
Adaptations to early-life adaptive UPR^ER^ in *C. elegans* depend on PEK-1 and ATFS-1 crosstalk. **A)** Lifespan analysis of *pek-1*(ok275) strain following treatment with TM. Kaplan–Meier survival plots of two independent experiments initiated with 100 animals/group, **** p ≤ 0.0001 Log-rank (Mantel-Cox) test. **B)** MitoTracker Red images of TM-treated day1 *pek-1*(ok275) worms. Scale 275μm. Mean +/- SEM; **** p ≤ 0.0001 Student’s t-test. **C)** Lifespan analysis of *atfs-1*(tm4525) following treatment with TM. Kaplan–Meier survival plots of two independent experiments initiated with 100 animals/group, **** p ≤ 0.0001 Log-rank (Mantel-Cox) test. **D)** MitoTracker Red images of TM-treated adult day1 *atfs-1*(tm4525) worms. Scale 275μm. Mean +/- SEM; **** p ≤ 0.0001 Student’s t-test. **E)** Lifespan assay of N2 Wild-type, *pek-1*(ok275), *atfs-1*(tm4525) and *pek-1*;*atfs-1* worms following TM treatment Kaplan– Meier survival plots of two independent experiments initiated with 100 animals per group. **F)** Mean lifespan of N2, *pek-1*(ok275), *atfs-1*(tm4525) and *pek-1*;*atfs-1* strains. * p ≤ 0.05, ** p ≤ 0.01, *** p ≤ 0.001, **** p ≤ 0.0001 One-way ANOVA. **G)** MitoTracker Red images of day1 N2, *pek-1*(ok275), *atfs-1* (tm4525) and *pek-1*;*atfs-1* strains. Data mean SEM of 45 animals/assay. *** p ≤ 0.001, **** p ≤ 0.0001 One-way ANOVA. **H)** Images of p*myo-3*::*mitogfp* reporter strain adult day4 N2 WT, *pek-1*(ok275), *atfs-1*(tm4525) and *pek-1*;*atfs-1* backgrounds. 3 independent experiments with 15 animals/condition, n=3; Scale 12μm. * p ≤ 0.05, ** p ≤ 0.01, *** p ≤ 0.001, **** p ≤ 0.0001 Chi-square. **I)** CeleST physical fitness parameters at day1 following TM treatment of N2 WT, *pek-1*, *atfs-1* and *pek-1*;*atfs-1* strains. Data from 3 independent experiments initiated with at least 10 animals per condition. Mean +/- SEM; ** p≤ 0.01, *** p ≤ 0.001 One-way ANOVA. **J)** CeleST physical fitness parameters that increase with age; average fitness parameters at day 1 following TM treatment of N2 WT, *pek-1*, *atfs-1* and *pek-1*;*atfs-1* mutant strains. Data from three independent experiments initiated with at least 10 animals/conditions. Mean +/- SEM; ** p ≤ 0.01 & **** p ≤ 0.0001 One-way ANOVA.

A double *pek-1*;*atfs-1* mutant was generated to clarify the possible interaction of these signalling pathways and potential crosstalk of the UPR^ER^ and UPR^mt^. The phenotypic effects of TM treatment in the N2 wild-type strain, *pek-1*, *atfs-1* and *pek-1*;*atfs-1* double mutant strains were determined. All mutant strains had decreased lifespan compared to the WT group which had increased longevity (Fig.7E&F, Suppl Table 1).). Interestingly, TM increased the size of N2 worms, but this effect was reversed in the *pek-1* mutant and absent in the *atfs-1* mutant (Suppl Fig.3E). The *pek-1;atfs-1* double mutant exhibited a more severe reduction in body size than either single mutant, both under basal conditions and following TM treatment (Suppl Fig.3E). All mutant strains treated with TM had decreased resistance to sodium arsenite and paraquat compared to N2 strain (Suppl. Fig.4E, Suppl Table 2&3).

Mitochondrial membrane potential was significantly reduced in both the *atfs-1* and *pek-1;atfs-1* mutants compared to N2, while no differences were observed between the untreated N2 and the *pek-1* mutant strain (Fig.7G). TM treatment only increased mitochondrial membrane potential in N2 WT worms, with all mutants displaying lower MitoTracker Red fluorescence intensity post-TM treatment (Fig.7G). To assess mitochondrial morphology, strains were crossed with the p*myo-3*::mitoGFP reporter strain and imaged at day4 of adulthood. Control N2 WT worms had a balanced mitochondrial network with approximately 20% filamentous, 40% intermediate, and 40% punctate mitochondria. The *pek-1* and *atfs-1* mutants demonstrated increased punctate or more fragmented mitochondrial network (Fig.7H). However, the *pek-1;atfs-1* double mutant displayed a highly disrupted mitochondrial network, with no filamentous mitochondria present. TM treatment increased the percentage of filamentous mitochondria in N2 WT worms (∼25%) and reduced the percentage of mitochondrial punctate (∼25%). In contrast, the mitochondrial morphology of *pek-1* and *atfs-1* single mutants remained unchanged after TM treatment, while the *pek-1;atfs-1* mutant exhibited further mitochondrial fragmentation (Fig.7H). Finally, to assess physiological activity, CeLeST analysis was performed. In untreated conditions, fitness-related parameters were similar between WT, *pek-1*, *atfs-1*, and *pek-1;atfs-1* mutants, while frailty parameters were slightly higher in the mutant strains (Fig.7I&J, Suppl Fig.3F). Following TM treatment, WT worms had increased wave initiation rate, travel speed and activity index indicating increased fitness, this was not observed in any of the mutant strains (Fig.7, Suppl Fig.3F).

Together, these results highlight the critical roles of PEK-1 and ATFS-1 in regulating lifespan, stress resistance and mitochondrial dynamics. The results indicate that under low levels of ER stress PEK-1 of UPR^ER^ establish a crosstalk with ATFS-1 of UPR^mt^ to enhance lifespan, healthspan, stress resistance and mitochondrial capacity. This demonstrates a functional interaction between these pathways previously unexplored in *C. elegans*.

In summary, the data presented using both an *in vitro* myogenesis model and *in vivo* whole organism *C. elegans*, reveals that adaptive UPR^ER^ signalling through PERK and UPR^mt^, induces the assembly of MERCS and regulates mitochondrial adaptations following a low dose of an ER stressor at an early developmental stage. These adaptations promote myogenesis and extension of lifespan by enhancing organelle turnover and mitochondrial function, demonstrating the essential crosstalk between ER and mitochondrial stress responses in maintaining cellular and organismal homeostasis under physiological stress conditions.

## Discussion

This study identified a pivotal role of adaptive UPR^ER^ signalling during early development, which regulates the assembly of MERCS, mitochondrial function, and subsequent physiological outcomes. Acute exposure to a low level of the ER stressor, TM, induced adaptive UPR^ER^ signalling and promoted adaptations in both *in vitr*o and *in vivo* models. The adaptative responses, including increased mitochondrial content and activity, that depend on the PERK arm of the UPR^ER^. A low concentration of TM enhanced myogenic potential in C2C12 myoblasts, while higher concentrations induced cell death. We propose that the improved myogenesis is a result of increased MERCS assembly and increased mitochondrial turnover and content. Similarly, in *C. elegans*, UPR^ER^ activation at an early developmental stage leads to significant lifespan extension and stress resistance during ageing. TM treatment promotes MERCS assembly, mitochondrial turnover and increased lipid metabolism in *C. elegans*. Previous studies in *C*. *elegans* have reported that an ER stress response can extend lifespan ^38^. Indeed, expression of a constitutively active neuronal XBP-1s can activate UPR^ER^ in distal tissues resulting in increased longevity ^39^, potentially by modulating lipid metabolism and increased lysosomal acidity ^33,40^. Mechanistically, the results presented here identified PEK-1 (PERK) as the essential regulator of adaptive UPR^ER^ signalling and determined crucial crosstalk between PEK-1 (UPR^ER^) and ATFS-1 (UPR^mt^) in maintaining mitochondrial integrity and promoting longevity. Although TM treatment increased longevity and preserved mitochondrial morphology, it was associated with decreased reproductive potential, supporting previous studies that have determined that UPR^ER^ induction promotes the allocation of organismal resources toward the maintenance of somatic cells and decreasing reproductive potential ^41^.

In both *in vitro* and *in vivo* models, TM treatment increased mitochondrial content, turnover and respiration. Disruption of the ER environment has been associated with downstream effects on mitochondrial function ^18^. The ER can coordinate mitochondrial dynamics by establishing contact sites through ER tubules ^23^. INF2 from the ER interacts with the OMM actin nucleator Spire1c to polymerise actin filaments around mitochondria ^42^, stimulating ER tubules to release Ca^2+^ ions into mitochondria through VDAC1, triggering the inner mitochondrial membrane to divide ^43^. MERCS also mediate the replication and distribution of mtDNA along the mitochondrial network during mitochondrial fission ^44,45^. ER tubules can also guide the position and timing of mitochondria fusion through tethering with mitochondria ^46^, as MERCS must be maintained to decrease mitochondrial motility ^47^. Narrow MERCS (<10nm) are characterised by an efficient transfer of molecules and signals, which enables a rapid response to metabolic changes and stress signals ^48^. It has been demonstrated that contact sites facilitate the efficient and rapid transfer of Ca^2+^ ions from the ER to mitochondria, crucial for maintaining cellular Ca^2+^ homeostasis and mitochondrial function ^49^. Contact sites also allow an efficient exchange of lipids between the ER and mitochondria, essential for membrane biosynthesis and the maintenance of mitochondrial structure ^50^. Long-distance MERCS (>30nm) limit the extent and speed of interactions, ensuring that only necessary signalling occurs ^48^. It has been reported that long-distance MERCS prevent excessive build-up of Ca^2+^ within the mitochondria, which can result in the opening of the mPTP, mitochondrial dysfunction and initiation of apoptotic signalling events ^49^.

The presence of ROS within MERCS has been reported to generate redox nanodomains between the two organelles, which have been proposed to mediate redox signalling and IP3R-mediated Ca^2+^ release via MERCS, resulting in the swelling of the mitochondrial matrix, reduction of the cristae and release of H_2_O_2_ ^51^. TEM analysis demonstrated an increase in MERCS following TM treatment at the embryonic stage which was maintained after the differentiation of myoblasts into myotubes and adult worms. The PERK arm of the UPR^ER^ can modulate mitochondrial capacity in response to acute ER stress, promoting the remodelling of the mitochondrial network by induction of SIMH ^18,19^. Adaptive UPR^ER^ signalling can induce the formation of MERCS, promoting Ca^2+^ transfer between these organelles and boosting mitochondrial metabolism by increasing the activity of the ETC and boosting the TCA cycle ^21^. PERK localises to mitochondrial-associated membranes (MAMs) and its interaction with the mitochondrial tethering protein MFN2 stabilises MERCS ^52^. Ablation of PERK disrupts ER morphology and reduces the number of MERCS, indicating that PERK is required at the ER for tethering mitochondrial membranes ^28^. How PERK regulates MERCS assembly is still uncertain. It has been demonstrated that cells lacking PERK have disrupted MERCS assembly ^2,28^. However, rescuing cells lacking PERK with kinase-dead PERK mutants restored MERCS assembly, indicating its kinase activity is not essential for maintaining MERCS ^2,28^. The stabilisation of MERCS depends on PERK’s cytosolic domain, as overexpression of PERK lacking this domain did not restore MERCS assembly ^28^, suggesting that PERK’s tethering role is structural rather than dependent on canonical PERK signalling ^28^. It has been suggested that the non-canonical role of PERK in regulating MERCS is via recruiting lipid transfer proteins such as Extended-Synaptotagmin-1 (E-Syt-1) ^53^. PERK also possess a conserved cysteine residue (Cys216) that can be reversibly oxidised, allowing the formation of covalent interactions with ERO1α and tightening of MERCS ^2^.

Activation of the UPR^ER^ during the transition from L4 larvae to adult day1 stage, had opposite effects compared to those reported following treatment at the embryonic stage, decreasing both lifespan and resistance to oxidative stress. In *C. elegans*, UPR^ER^ activation in response to ER stress begins to decline relatively early in the ageing process ^39,54^, coinciding with the peak of the reproductive period ^55^. It has been reported that UPR^ER^ induction promotes the allocation of organismal resources toward the maintenance of somatic cells, thereby decreasing reproductive potential ^41^. As cells age, their ability to maintain proteostasis diminishes, highlighting the importance of sustaining UPR^ER^ function to prevent age-related declines in homeostasis ^41^. Transcriptomic analysis revealed that TM treatment during early development upregulated genes involved in lipid metabolism and autophagy, contributing to lifespan extension. In contrast, TM treatment in later life reduced lipid metabolism, impairing survival. Enhanced longevity in *C*. *elegans* is linked to the transcriptional upregulation of genes related to lysosomal function and the modulation of fatty acid desaturases, lysosomal lipases and general lipophagy ^33,40,56^. Lipid staining confirmed these changes, with embryo TM-treated worms having lower lipid stores and increased expression of genes involved in lipid synthesis and decreased lysosomal pH, while adult TM-treated worms had higher lipid levels and decreased expression of genes involved in autophagy. Lipid biosynthesis requires phospholipid exchange between the ER and the mitochondria at MERCS. Phosphatidylserine at the ER membrane can transfer to mitochondria, where it is converted into phosphatidylethanolamine and then transported back to the ER to form phosphatidylcholine ^57^. The data supports recent studies on the role of PERK in promoting lipid trafficking at MERCS ^53^.

To investigate the IRE-1 arm of the UPR^ER^, mutant strains for *ire-1* and *xbp-1* were used, but all larvae arrested following TM treatment. Although the mutant strains of the IRE-1 arm were viable, when faced with an additional ER stress during early development, the larvae arrested. A similar effect has been reported in *ire-1* or *xbp-1* mutant strains with the deletion of *atf-6* or *pek-1* ^37^. In this study, *pek-1* mutants subjected to TM treatment exhibited a reduction in lifespan, oxidative stress resistance and mitochondrial membrane potential ^37^. In myoblasts, PERK inhibition blocked the myogenic and mitochondrial adaptations. Cells deficient in PERK, exhibit disrupted ETC activity and increased mitochondrial ROS ^58^. The *atfs-1* mutant strain had a similar phenotype to the *pek-1* mutant following TM treatment, which had an opposite effect in both mutant strains compared to the N2 wild-type strain. The observed phenotypic changes were also reduced following TM treatment in *pek-1;atfs-1* double mutants, suggesting that *pek-1* and *atfs-1* operate within the same signalling pathway. Based on the results presented and previous studies, PERK can determine mitochondrial remodelling through the assembly of MERCS, possibly via alterations in the redox environment and recruitment of lipid transfer proteins at MAMs. Activation of UPR^mt^ following ER stress aims to restore mitochondrial function and proteostasis by promoting mitochondrial repair and mitophagy to prevent mitochondrial dysfunction ^59^.

In summary, the results highlight that activation of an adaptive UPR^ER^ response during early developmental stages promoted PERK-dependent assembly of MERCS that determines mitochondrial function and dynamics, as well as the regulation of genes related to lysosomal function and lipid metabolism. The interaction between PEK-1 (UPR^ER^) and ATFS-1 (UPR^mt^) in response to early-life ER stress was essential for maintaining mitochondrial dynamics and function, lifespan extension and stress resistance.

## SUPPLEMENTAL INFORMATION

**Suppl. Figure 1:**
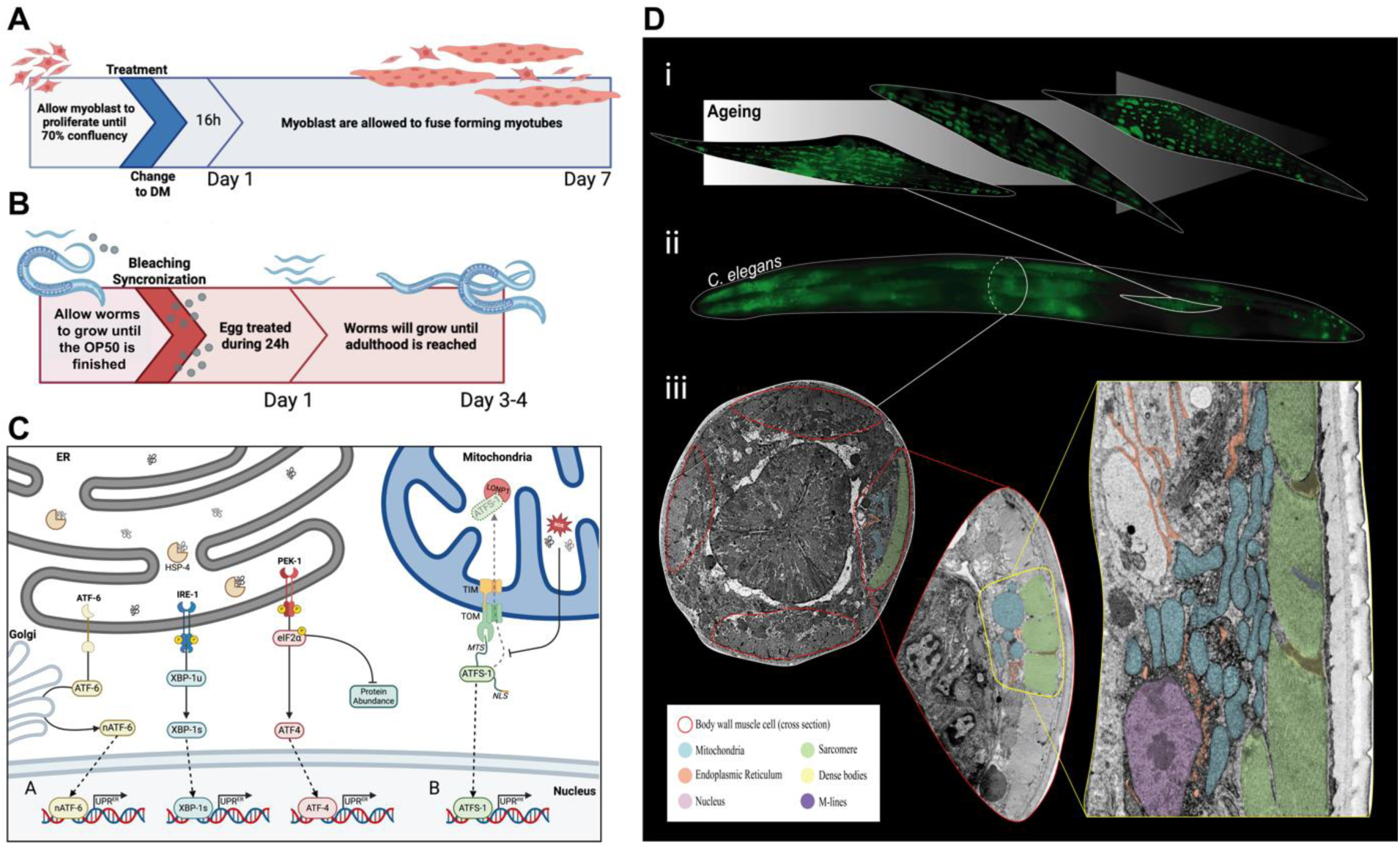
Myoblast and *C. elegans* models for determination of the effects of adaptive UPR^ER^ signalling during early developmental stages. **A)** Schematic diagram of the TM treatment protocol for C2C12 myoblasts. **B)** Schematic diagram of the TM treatment protocol of *C. elegans* at embryo stage. **C)** The UPR^ER^ and UPR^mt^ of *C. elegans*. **D)** Organelle ultrastructure from the body wall muscle of *C. elegans*. (i) Muscle cell ageing visualised using a p*myo-3*::mitoGFP translational fusion highlighting the mitochondrial network in body wall cells from a young day 2, 4 & 6 adult demonstrating mitochondrial network disorganisation. (ii) The organisation of body wall muscle cells in a mature hermaphrodite *C. elegans* (seen in a dorsal oblique view) was observed in a genetically modified worm expressing the P*myo-3*::mitoGFP reporter transgene. (iii) TEM cross-section images of body wall muscle cells. The first image shows the distribution of the four muscle quadrants (containing two rows of cells in each) from the pharynx area. The second image demonstrates a cross-section of the body wall cell, in which mitochondria, ER and Sarcomeres can be distinguished. High-resolution TEM section of the nucleus, mitochondria, ER and sarcomeres (with dense bodies and M-lines) in the body wall muscle cell.

**Suppl. Figure 2:**
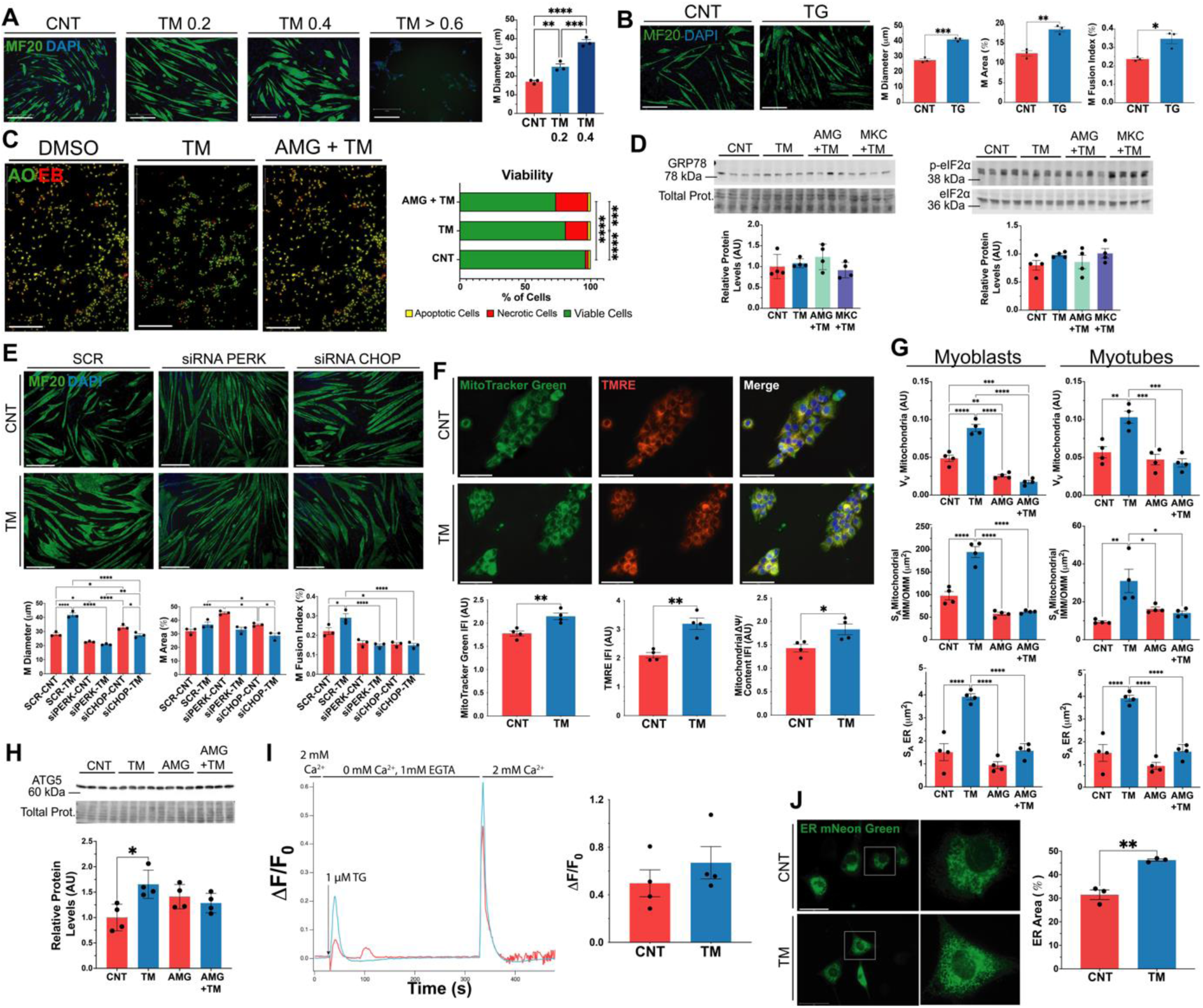
Tunicamycin treatment effect on myoblasts including myogenesis, mitochondrial and ER content and volume. A) C2C12 myoblasts were exposed to TM 0.2, 0.4, 0.6, 0.8 and 1 μg/ml for 8h. Media was replaced with differentiation media and cells were allowed to differentiate for 7 days. MF20 immunostaining demonstrated that TM treatment improved the myogenic potential of the myoblast population compared to the control; this effect is lost when the concentration of TM exceeds the 0.4 μg/ml. ** p ≤ 0.01, *** p ≤ 0.001 & **** p ≤ 0.0001 One-way ANOVA. B) C2C12 myoblasts were exposed to thapsigargin (TG) 10 nM for 8h. Media was replaced with differentiation media and cells were allowed to differentiate for 7 days. MF20 immunostaining demonstrated that TG treatment significantly increases myotube diameter, area and fusion index. * p ≤ 0.05, ** p ≤ 0.01 & *** p ≤ 0.001 Student’s t-test. C) C2C12 myoblasts were exposed to TM 0.2μg/ml and AMG PERK44 (PERK inhibitor) 2μM for 8h. Media was replaced with differentiation media and cells were allowed to recover for 16h. Treatment with TM resulted in a significant increase in cell death compared to the control group; this effect is exacerbated with the inclusion of the PERK inhibitor. *** p ≤ 0.001 & **** p ≤ 0.0001 Chi-square. D) Protein expression levels of GRP78 and p-EIF2α/EIF2α assessed by Western Blot, normalised against total protein (Ponceau staining). One-way ANOVA. E) C2C12 myoblasts were exposed to TM 0.2 μg/ml or, alongside Scr, siRNA for PERK or siRNA for CHOP during 8h. Media was replaced with differentiation media, and cells allowed to differentiate for 7 days. MF20 immunostaining demonstrated that the TM treatment increased the myogenic potential of the myoblast population compared to the control, but this effect is lost with the silencing of RNA for PERK and CHOP. * p ≤ 0.05, ** p ≤ 0.01, *** p ≤ 0.001 & **** p ≤ 0.0001 One-way ANOVA. F) TM treatment resulted in a significant increase in MitoTracker green fluorescence intensity. TM treatment increased TMRE staining, indicating an increase in mitochondrial membrane potential. The TMRE:MitoTracker green ratio increased in the TM group,. Representative images; n=4. * p ≤ 0.05, ** p ≤ 0.01, *** p ≤ 0.001 & **** p ≤ 0.0001 Student’s t-test. G) Mitochondria volume fraction (V_V_), IMM/OMM ratio (S_A_) and Surface area (S_A_) of ER membrane and mitochondria calculated from TEM sections from a population of myoblasts and myotubes. * p ≤ 0.05, ** p ≤ 0.01, *** p ≤ 0.001 & **** p ≤ 0.0001 One-way ANOVA. H) Protein expression levels of ATG5 were assessed by Western Blot, and normalised against total protein (Ponceau staining);. * p ≤ 0.05 One-way ANOVA. I) Amplitude values of [Ca^2+^]i increase show that the TM treated myoblasts exhibited a slightly augmented Store-Operated-Calcium-Entry (SOCE) compared to control. Student’s t-test. J) The area fraction of the ER surface against the total area of the cells. Representative images; green: ER mNeon Green, blue: DAPI; n=3. Scale 275 μm. ** p ≤ 0.01 Student’s t-test.

**Suppl. Figure 3:**
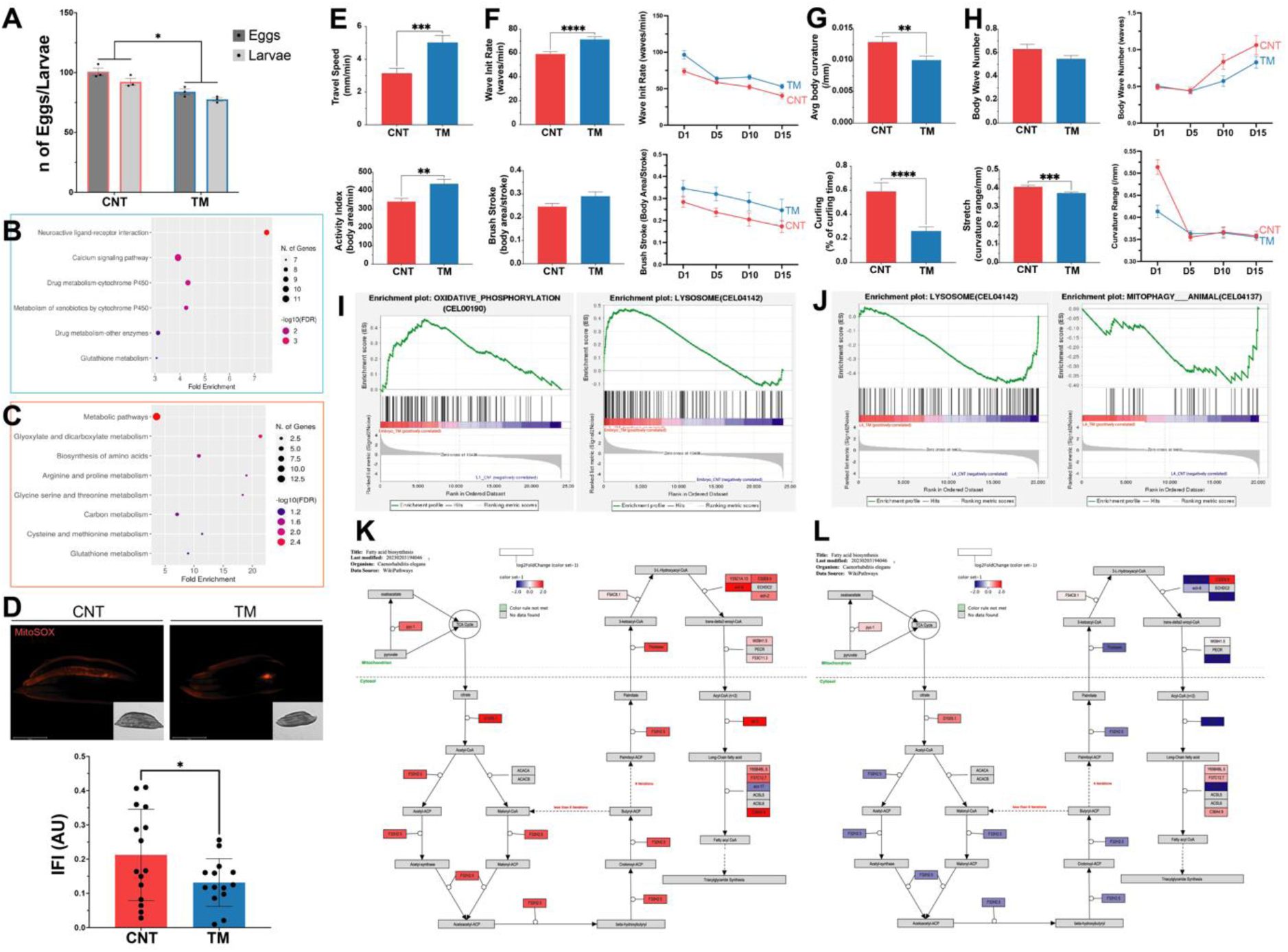
Early-life Tunicamycin treatment of *C. elegans* promotes cellular and physiological adaptations. **A)** Number of laid embryos compared to number of viable larvae in N2 worms treated with TM. * p ≤ 0.05 One-way ANOVA. **B)** Dot plot of Shiny GO KEGG analysis of EMBRYO DEGs. Significantly downregulated pathways’ dotplot outlined in blue. **C)** Dot plot of Shiny GO KEGG analysis of L4 DEGs. Significantly upregulated pathways’ dotplot outlined in red. **D)** N2 Wild-type worms incubated with MitoSOX dye, an indicator of mitochondrial ROS production. Representative images of three independent experiments initiated with 15 animals/ strain; red: MitoSOX. Scale 275μm. Error bars SEM; * p ≤ 0.05 Student’s t-test. **E & F)** CeleST physical fitness parameters that decrease with age; average fitness parameters of days 1, 5, 10, & 15. Mean +/- SEM; **** p ≤ 0.0001 Student’s t-test. **G & H)** CeleST physical fitness parameters that increase with age; average fitness parameters of days 1, 5, 10, & 15. Mean +/- SEM; *** p ≤ 0.001 Student’s t-test. **I & J)** GSEA pathways that are significantly enriched following treatment of embryos (I) or L4 worms (J). K) PathVisio pathway analysis of EMBRYO DEGs revealed an upregulation of proteins involved in fatty acid biosynthesis in worms treated with TM. L) PathVisio pathway analysis of L4 DEGs revealed the downregulation of proteins involved in fatty acid biosynthesis in worms treated with TM.

**Suppl. Figure 4:**
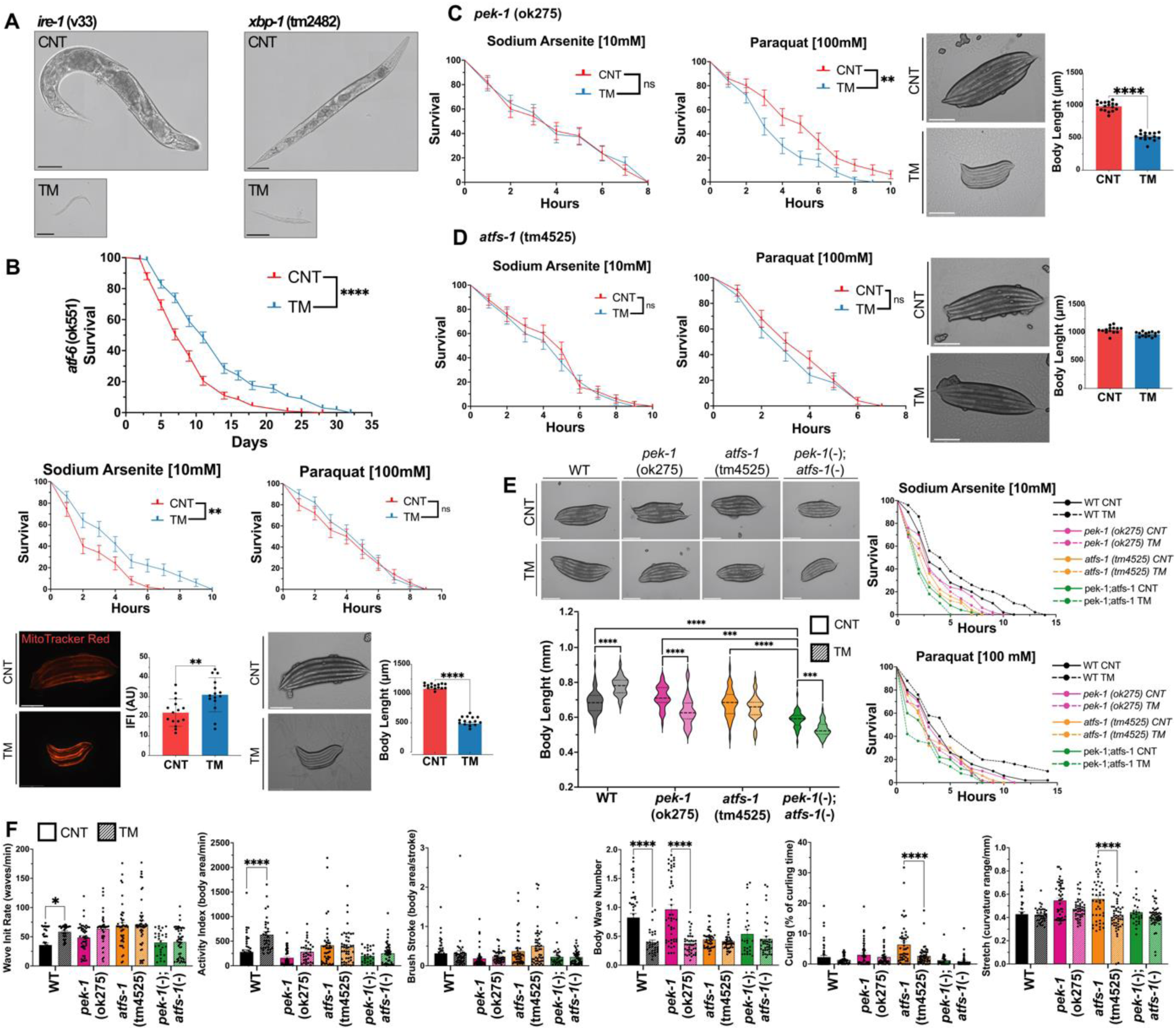
Effect of the UPR^ER^ in the adaptations to early-life TM treatment of *C. elegans*. **A)** Representative transmitted light images of TM treated *ire-1* (v33) worms and *xbp-1*(tm2482) worms. **B)** TM treatment of *atf-6* (ok551) increased lifespan at 20°C. Kaplan–Meier survival plots of two independent experiments initiated with 100 animals per group, Log-rank (Mantel-Cox) test, **** p ≤ 0.0001. Increased survival of TM-treated *atf-6*(ok551) in 10 mM Sodium Arsenite and no effect on survival in 100 mM Paraquat compared to control. Kaplan–Meier survival plots of two independent experiments initiated with 50 animals per group by Log-rank (Mantel-Cox) test, ** p ≤ 0.01. Representative MitoTracker Red staining of adult day 1 *atf-6*(ok551) worms treated with TM. Scale 275 μm. Mean +/- SEM; ** p ≤ 0.01 Student’s t-test. Representative transmitted light images of adult day 1 *atf-6*(ok551) mutant worms. Scale 275μm. Error bars SEM; ** p ≤ 0.01 Student’s t-test. **C)** TM treatment of *pek-1*(ok275) did not affect survival in 10 mM Sodium Arsenite and decreased survival in 100 mM Paraquat compared to control. Kaplan–Meier survival plots of two independent experiments were initiated with 50 animals per group using log-rank (Mantel-Cox) test, ** p ≤ 0.01. Representative transmitted light images of adult day 1 *pek-1*(ok275) worms. Scale 275 μm. Mean +/- SEM; **** p ≤ 0.0001 Student’s t-test. **D)** TM treatment of *atfs-1*(tm4525) had no effect on survival in 10 mM Sodium Arsenite and 100 mM Paraquat compared to controls. Kaplan–Meier survival plots of two independent experiments initiated with 50 animals per group by Log-rank (Mantel-Cox) test. Representative transmitted light images of adult day 1 *atfs-1*(tm4525) worms. Scale 275μm. Mean +/- SEM; Student’s t-test. **E)** Body length of adult day 1 worms following TM treatment in N2 *pek-1*(ok275), *atfs-1*(tm4525) and *pek-1;atfs-1* strains. Mean +/- SEM of 45 animals per assay. *** p ≤ 0.001 & **** p ≤ 0.0001 One-way ANOVA. **F)** CeleST physical fitness parameters that change with age; average fitness parameters of day 1 TM-treated N2, *pek-1*, *atfs-1*, and *pek-1;atfs-1* mutant strains. Data from three independent experiments initiated with at least 10 animals per condition. Scale 275μm. Mean +/- SEM; * p ≤ 0.05 & **** p ≤ 0.0001 One-way ANOVA.

**Suppl. Table 1:**
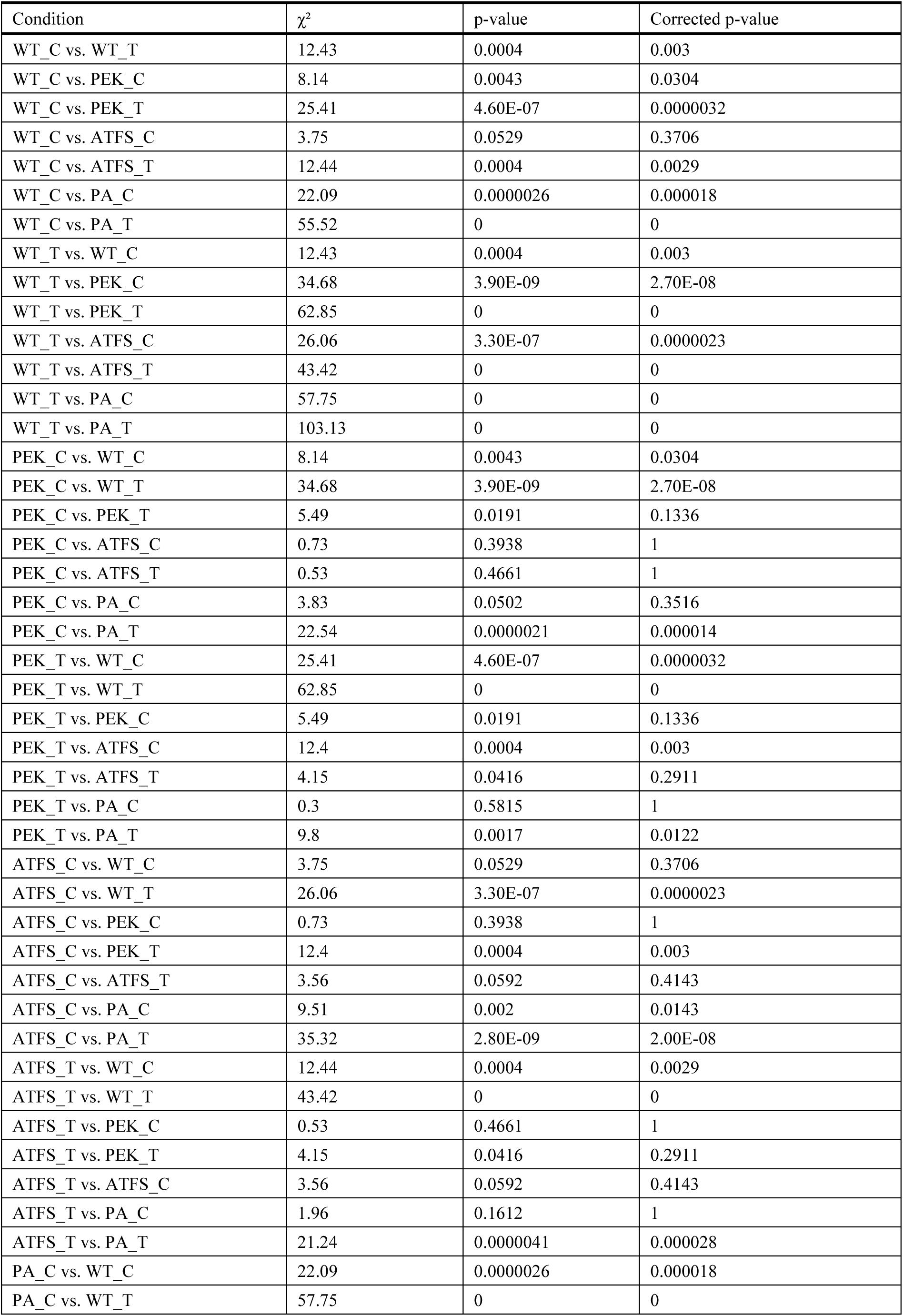
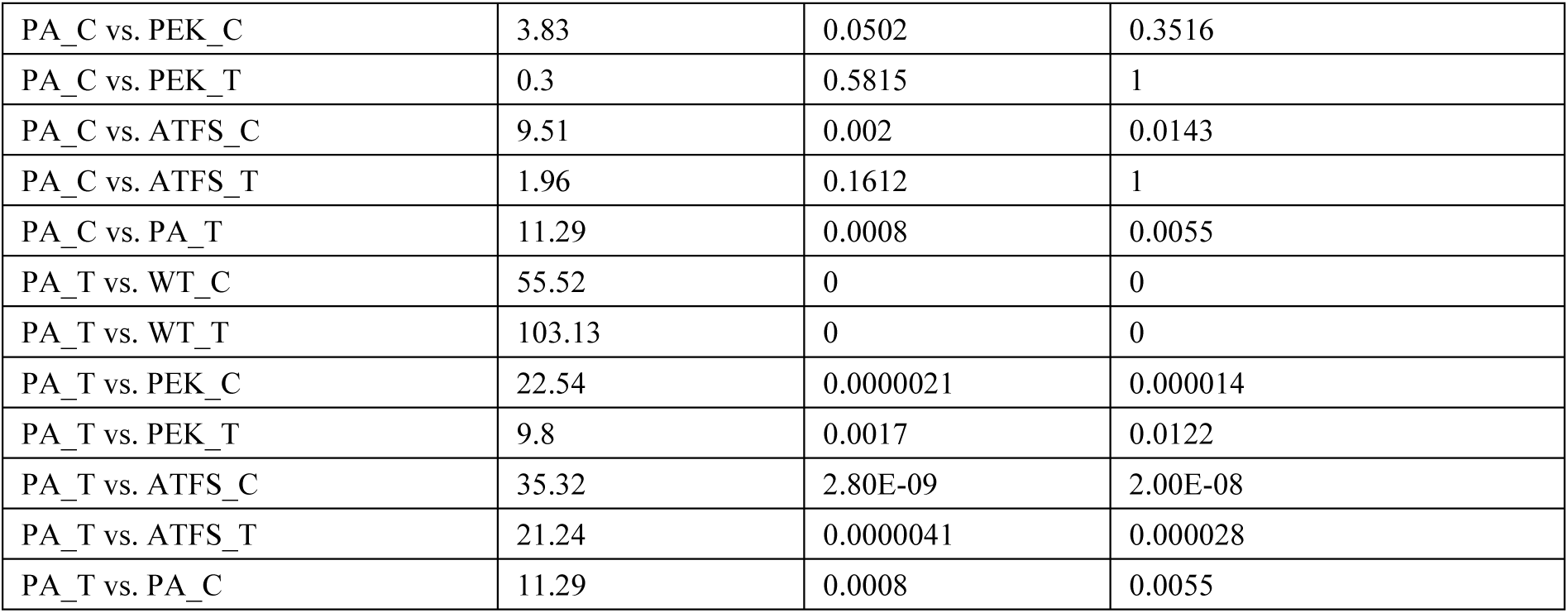
Multiple comparison lifespan analysis at 20°C of wild-type (WT), *pek-1*(ok275) (PEK), *atfs-1(*tm4525) (ATFS) and *pek-1;atfs-1*. Worms were treated with 1.25 μg/ml TM at EMBRYO stage during 24h, two independent experiments initiated with 100 animals per group.

**Suppl. Table 2:** Multiple comparison Sodium Arsenite survival analysis at 20°C of wild-type (N2), *pek-1*(ok275) (PEK), *atfs-1*(tm4525) (ATFS) and pek-1;atfs-1. Worms were treated with 1.25 μg/ml TM EMBRYO stage during 24h, two independent experiments initiated with 50 animals per group.

| Condition | $\chi^2$ | p-value | Corrected p-value |
| --- | --- | --- | --- |
| WT_C vs. WT_T | 4.48 | 0.0343 | 0.2402 |
| WT_C vs. PEK_C | 1.59 | 0.2067 | 1 |
| WT_C vs. PEK_T | 5.09 | 2.41E-02 | 0.1685 |
| WT_C vs. ATFS_C | 6.57 | 0.0104 | 0.0727 |
| WT_C vs. ATFS_T | 10.73 | 0.0011 | 0.0074 |
| WT_C vs. PA_C | 14.43 | 0.0001 | 0.001 |
| WT_C vs. PA_T | 21.66 | 0.0000032 | 0.000023 |
| WT_T vs. WT_C | 4.48 | 0.0343 | 0.2402 |
| WT_T vs. PEK_C | 8.27 | 4.00E-03 | 2.82E-02 |
| WT_T vs. PEK_T | 13.18 | 0.0003 | 0.002 |
| WT_T vs. ATFS_C | 15.31 | 1.00E-04 | 0.0006 |
| WT_T vs. ATFS_T | 20.87 | 0.0000049 | 0.000034 |
| WT_T vs. PA_C | 26.91 | 2.10E-07 | 0.0000015 |
| WT_T vs. PA_T | 36.73 | 0 | 0 |
| PEK_C vs. WT_C | 1.59 | 0.2067 | 1 |
| PEK_C vs. WT_T | 8.27 | 4.00E-03 | 2.82E-02 |
| PEK_C vs. PEK_T | 1.46 | 0.2267 | 1 |
| PEK_C vs. ATFS_C | 2.77 | 0.0961 | 0.6726 |
| PEK_C vs. ATFS_T | 6.2 | 0.0128 | 0.0897 |
| PEK_C vs. PA_C | 8.27 | 0.004 | 0.0283 |
| PEK_C vs. PA_T | 13.96 | 0.0002 | 0.0013 |
| PEK_T vs. WT_C | 5.09 | 2.41E-02 | 0.1685 |
| PEK_T vs. WT_T | 13.18 | 0.0003 | 0.002 |
| PEK_T vs. PEK_C | 1.46 | 0.2267 | 1 |
| PEK_T vs. ATFS_C | 0.27 | 0.6021 | 1 |
| PEK_T vs. ATFS_T | 1.86 | 0.1721 | 1 |
| PEK_T vs. PA_C | 3.23 | 0.0722 | 0.5057 |
| PEK_T vs. PA_T | 8.5 | 0.0035 | 0.0248 |
| ATFS_C vs. WT_C | 6.57 | 0.0104 | 0.0727 |
| ATFS_C vs. WT_T | 15.31 | 1.00E-04 | 0.0006 |
| ATFS_C vs. PEK_C | 2.77 | 0.0961 | 0.6726 |
| ATFS_C vs. PEK_T | 0.27 | 0.6021 | 1 |
| ATFS_C vs. ATFS_T | 1 | 0.3165 | 1 |
| ATFS_C vs. PA_C | 2.3 | 0.1296 | 0.9072 |
| ATFS_C vs. PA_T | 7.55 | 6.00E-03 | 4.20E-02 |
| ATFS_T vs. WT_C | 10.73 | 0.0011 | 0.0074 |
| ATFS_T vs. WT_T | 20.87 | 0.0000049 | 0.000034 |
| ATFS_T vs. PEK_C | 6.2 | 0.0128 | 0.0897 |
| ATFS_T vs. PEK_T | 1.86 | 0.1721 | 1 |
| ATFS_T vs. ATFS_C | 1 | 0.3165 | 1 |
| ATFS_T vs. PA_C | 0.32 | 0.5729 | 1 |
| ATFS_T vs. PA_T | 3.26 | 0.0708 | 0.4957 |
| PA_C vs. WT_C | 14.43 | 0.0001 | 0.001 |
| PA_C vs. WT_T | 26.91 | 2.10E-07 | 0.0000015 |
| PA_C vs. PEK_C | 8.27 | 0.004 | 0.0283 |
| PA_C vs. PEK_T | 3.23 | 0.0722 | 0.5057 |
| PA_C vs. ATFS_C | 2.3 | 0.1296 | 0.9072 |
| PA_C vs. ATFS_T | 0.32 | 0.5729 | 1 |
| PA_C vs. PA_T | 1.63 | 0.2014 | 1 |
| PA_T vs. WT_C | 21.66 | 0.0000032 | 0.000023 |
| PA_T vs. WT_T | 36.73 | 0 | 0 |
| PA_T vs. PEK_C | 13.96 | 0.0002 | 0.0013 |
| PA_T vs. PEK_T | 8.5 | 0.0035 | 0.0248 |
| PA_T vs. ATFS_C | 7.55 | 6.00E-03 | 4.20E-02 |
| PA_T vs. ATFS_T | 3.26 | 0.0708 | 0.4957 |
| PA_T vs. PA_C | 1.63 | 0.2014 | 1 |

**Suppl. Table 3:** Multiple comparison Paraquat survival analysis at 20°C of N2 wild-type (WT), *pek-1*(ok275) (PEK), *atfs-1(*tm4525) (ATFS) and *pek-1*;*atfs-1* strains. Worms were treated with 1.25 μg/ml TM EMBRYO stage during 24h, two independent experiments initiated with 50 animals per group.

| Condition | $\chi^2$ | p-value | Corrected p-value |
| --- | --- | --- | --- |
| WT_C vs. WT_T | 4.13 | 0.0422 | 0.2956 |
| WT_C vs. PEK_C | 0.55 | 0.4593 | 1 |
| WT_C vs. PEK_T | 3.58 | 5.84E-02 | 0.4088 |
| WT_C vs. ATFS_C | 1.97 | 0.1606 | 1 |
| WT_C vs. ATFS_T | 2.98 | 0.0846 | 0.5919 |
| WT_C vs. PA_C | 5.34 | 0.0208 | 0.1456 |
| WT_C vs. PA_T | 10.29 | 0.0013 | 0.0094 |
| WT_T vs. WT_C | 4.13 | 0.0422 | 0.2956 |
| WT_T vs. PEK_C | 7.14 | 7.50E-03 | 5.27E-02 |
| WT_T vs. PEK_T | 11.47 | 0.0007 | 0.0049 |
| WT_T vs. ATFS_C | 9.42 | 2.10E-03 | 0.015 |
| WT_T vs. ATFS_T | 10.53 | 0.0012 | 0.0082 |
| WT_T vs. PA_C | 13.53 | 0.0002 | 0.0016 |
| WT_T vs. PA_T | 21.12 | 0.0000043 | 0.00003 |
| PEK_C vs. WT_C | 0.55 | 0.4593 | 1 |
| PEK_C vs. WT_T | 7.14 | 7.50E-03 | 5.27E-02 |
| PEK_C vs. PEK_T | 1.72 | 0.1903 | 1 |
| PEK_C vs. ATFS_C | 0.77 | 0.3815 | 1 |
| PEK_C vs. ATFS_T | 1.47 | 0.2246 | 1 |
| PEK_C vs. PA_C | 2.71 | 0.0999 | 0.699 |
| PEK_C vs. PA_T | 6.92 | 0.0085 | 0.0595 |
| PEK_T vs. WT_C | 3.58 | 5.84E-02 | 0.4088 |
| PEK_T vs. WT_T | 11.47 | 0.0007 | 0.0049 |
| PEK_T vs. PEK_C | 1.72 | 0.1903 | 1 |
| PEK_T vs. ATFS_C | 0.37 | 0.5422 | 1 |
| PEK_T vs. ATFS_T | 0.08 | 0.7726 | 1 |
| PEK_T vs. PA_C | 0.22 | 0.6361 | 1 |
| PEK_T vs. PA_T | 2.56 | 0.1095 | 0.7668 |
| ATFS_C vs. WT_C | 1.97 | 0.1606 | 1 |
| ATFS_C vs. WT_T | 9.42 | 2.10E-03 | 0.015 |
| ATFS_C vs. PEK_C | 0.77 | 0.3815 | 1 |
| ATFS_C vs. PEK_T | 0.37 | 0.5422 | 1 |
| ATFS_C vs. ATFS_T | 0.1 | 0.754 | 1 |
| ATFS_C vs. PA_C | 1.12 | 0.2905 | 1 |
| ATFS_C vs. PA_T | 4.65 | 3.10E-02 | 2.17E-01 |
| ATFS_T vs. WT_C | 2.98 | 0.0846 | 0.5919 |
| ATFS_T vs. WT_T | 10.53 | 0.0012 | 0.0082 |
| ATFS_T vs. PEK_C | 1.47 | 0.2246 | 1 |
| ATFS_T vs. PEK_T | 0.08 | 0.7726 | 1 |
| ATFS_T vs. ATFS_C | 0.1 | 0.754 | 1 |
| ATFS_T vs. PA_C | 0.7 | 0.4024 | 1 |
| ATFS_T vs. PA_T | 4.1 | 0.043 | 0.3008 |
| PA_C vs. WT_C | 5.34 | 0.0208 | 0.1456 |
| PA_C vs. WT_T | 13.53 | 0.0002 | 0.0016 |
| PA_C vs. PEK_C | 2.71 | 0.0999 | 0.699 |
| PA_C vs. PEK_T | 0.22 | 0.6361 | 1 |
| PA_C vs. ATFS_C | 1.12 | 0.2905 | 1 |
| PA_C vs. ATFS_T | 0.7 | 0.4024 | 1 |
| PA_C vs. PA_T | 1.16 | 0.282 | 1 |
| PA_T vs. WT_C | 10.29 | 0.0013 | 0.0094 |
| PA_T vs. WT_T | 21.12 | 0.0000043 | 0.00003 |
| PA_T vs. PEK_C | 6.92 | 0.0085 | 0.0595 |
| PA_T vs. PEK_T | 2.56 | 0.1095 | 0.7668 |
| PA_T vs. ATFS_C | 4.65 | 3.10E-02 | 2.17E-01 |
| PA_T vs. ATFS_T | 4.1 | 0.043 | 0.3008 |
| PA_T vs. PA_C | 1.16 | 0.282 | 1 |

## ACKNOWLEDGMENTS

We would like to sincerely thank the Tavernarakis lab (University of Crete) for providing the IR2539 mitophagy reporter strain and the facilities and scientific and technical assistance of the Anatomy Imaging and Microscopy Facility at the University of Galway (https://imaging.universityofgalway.ie/imaging/) for electron microscopy work. JCCM studentship is funded by a Hardiman Scholarship and the College of Nursing Medicine and Health Sciences, University of Galway. QX (202006370047) and PL (202206370063) studentships are funded by the Chinese Scholarship Council (CSC).

## AUTHOR CONTRIBUTIONS

**José C. Casas-Martinez:** Writing – review & editing, Writing – original draft, Visualisation, Methodology, Formal analysis, Conceptualisation. **Qin Xia:** Methodology, Formal analysis. **Penglin Li:** Methodology, Formal analysis. **Antonio Miranda-Vizuete:** Writing – review & editing, Resources, Methodology. **Emma McDermott:** Resources, Methodology. **Peter Dockery:** Writing – review & editing, Resources, Methodology, Formal analysis. **Leo R. Quinlan**: Writing – review & editing, Resources, Methodology **Katarzyna Goljanek-Whysall:** Writing – review & editing, Resources, Methodology. **Afshin Samali:** Writing – review & editing, Supervision, Resources. **Brian McDonagh:** Writing – review & editing, Writing – original draft, Supervision, Resources, Methodology, Formal analysis, Conceptualisation

## STAR METHODS

### KEY RESOURCES TABLE

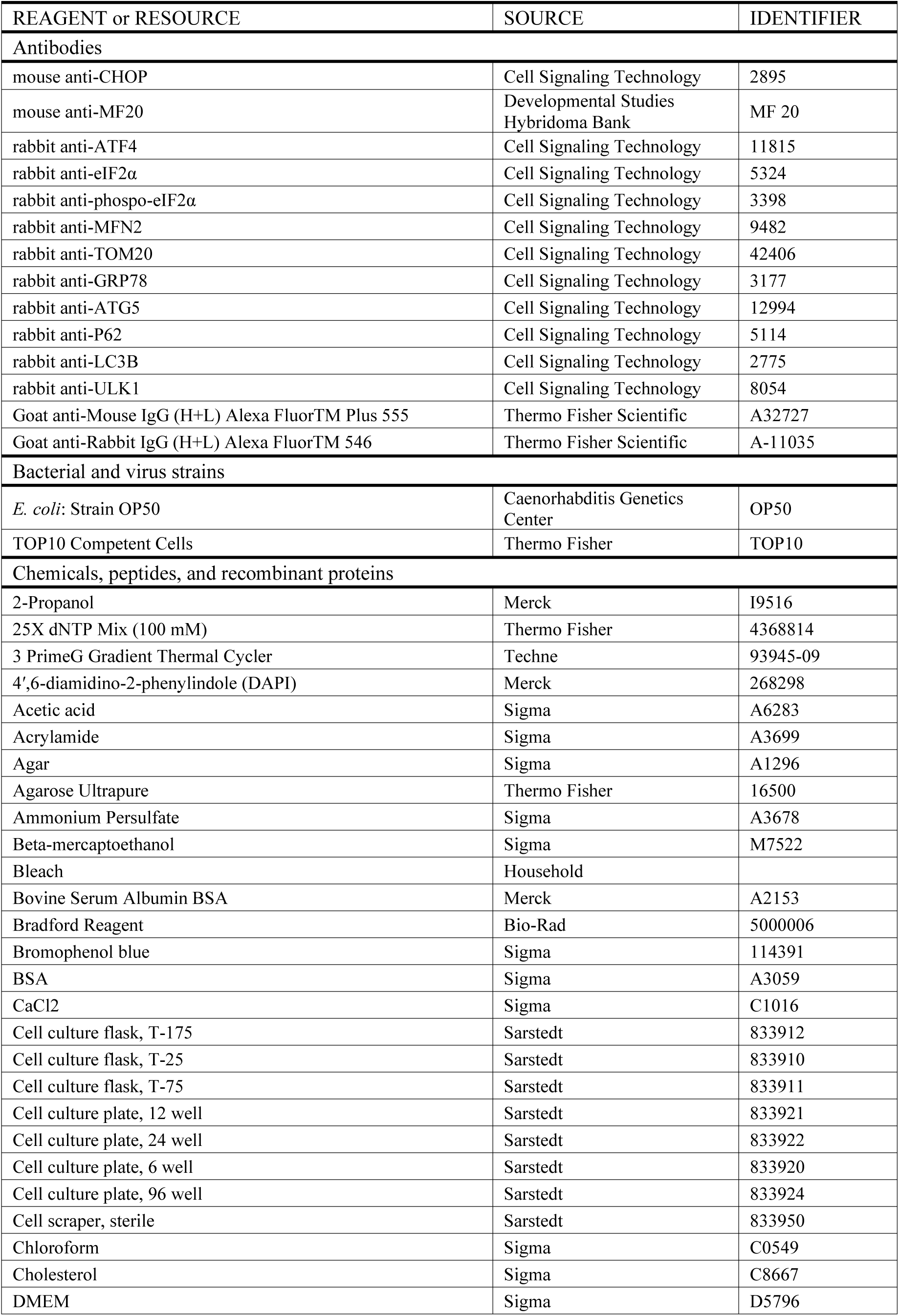
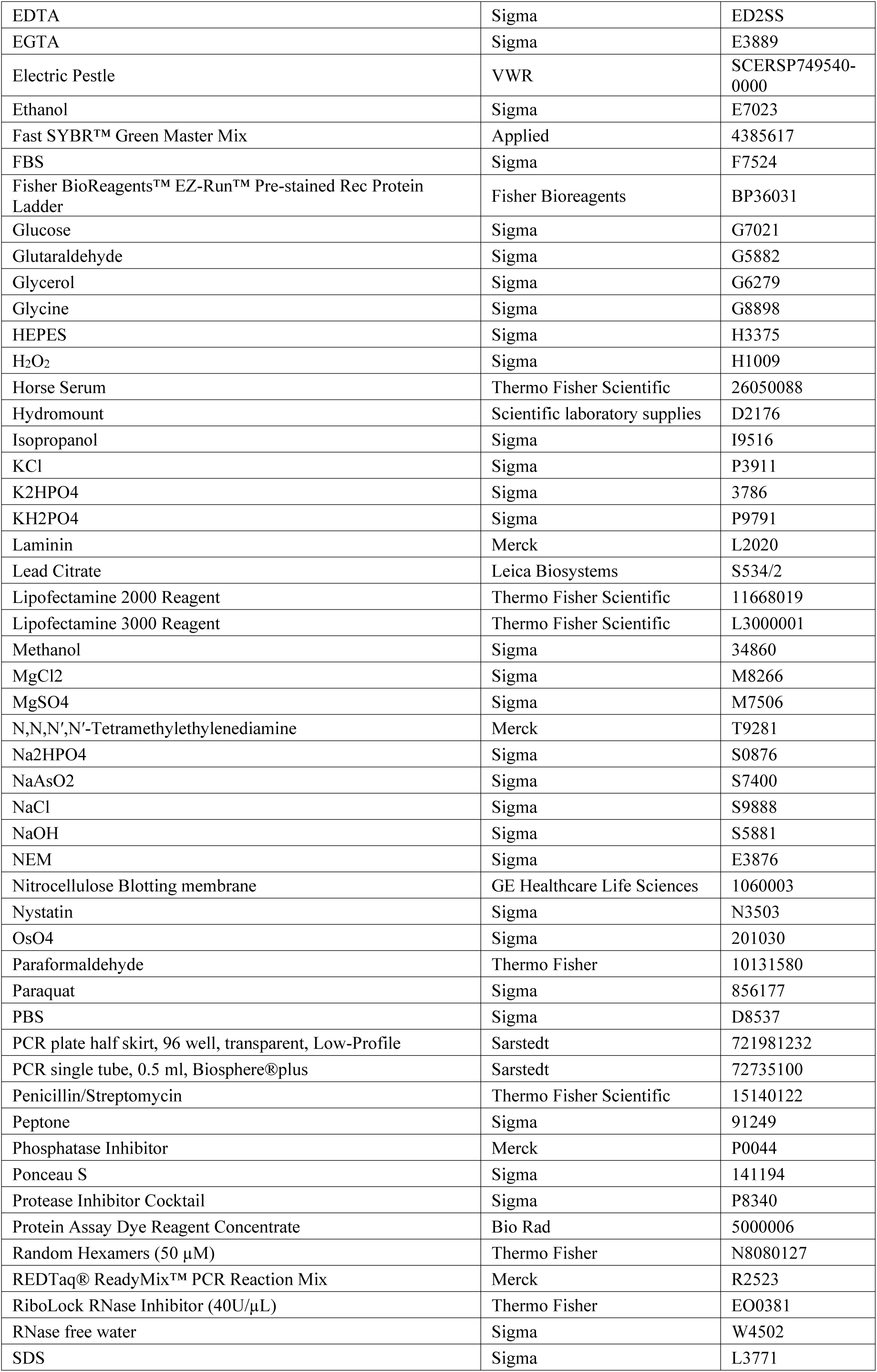
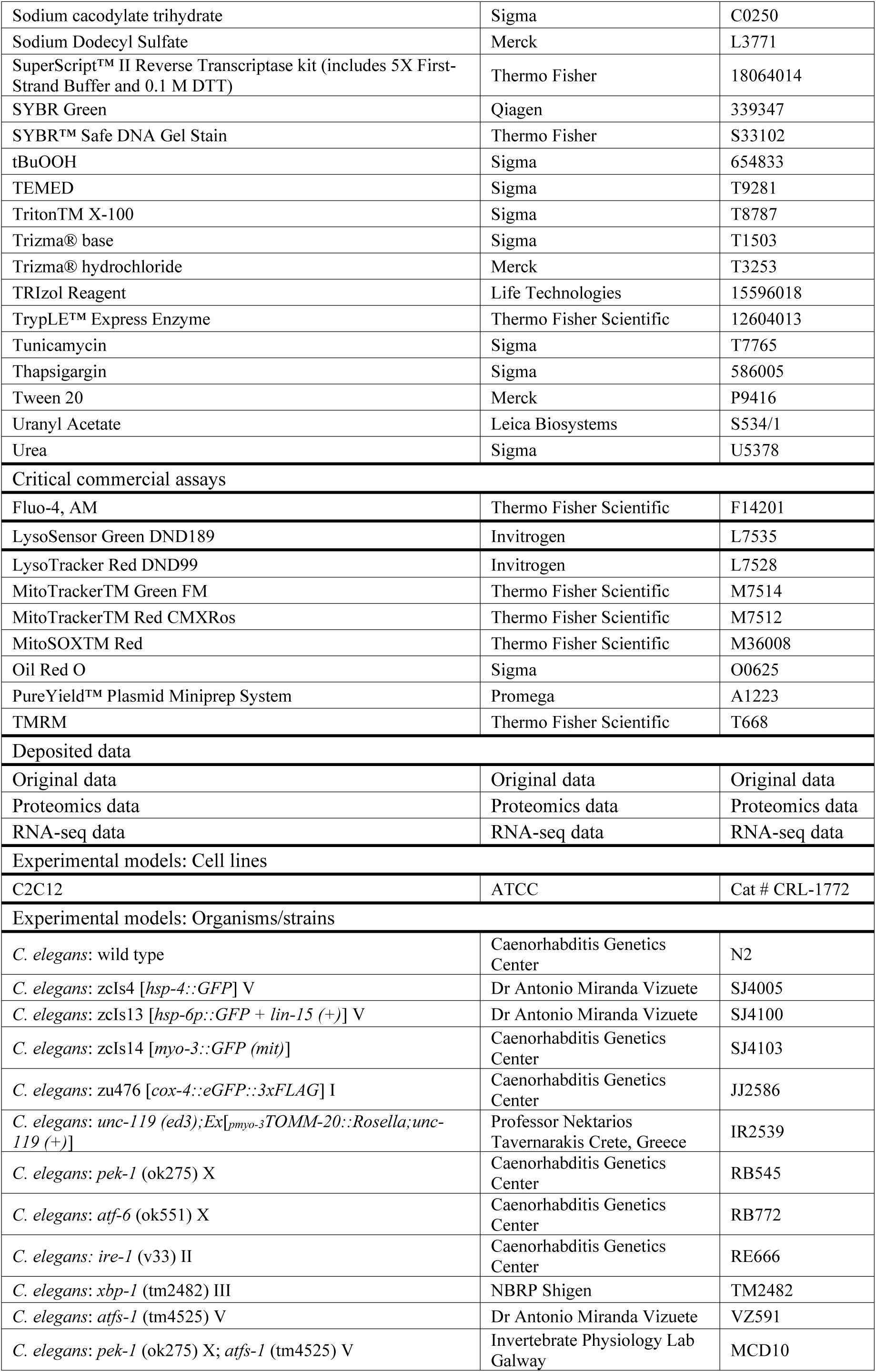
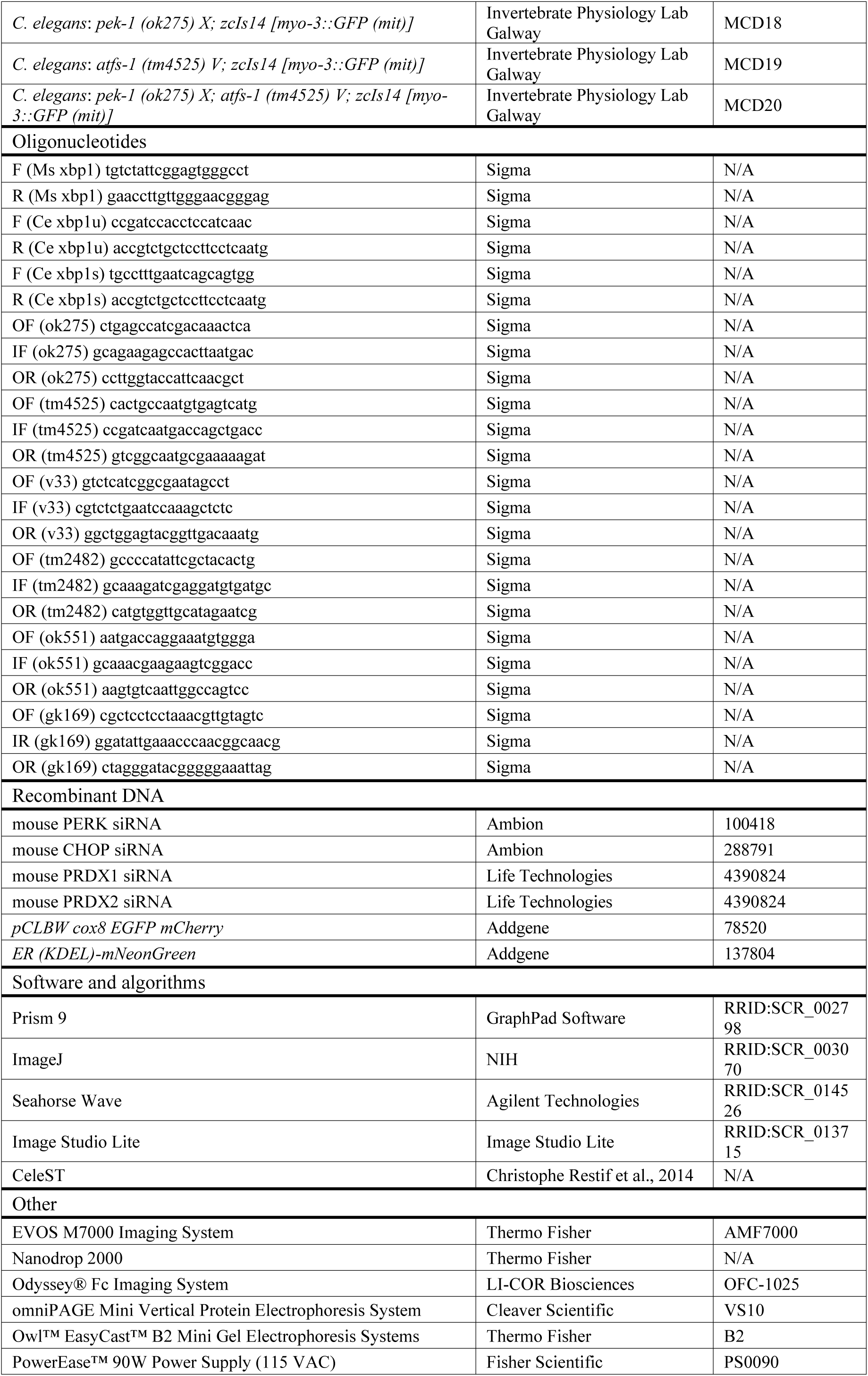
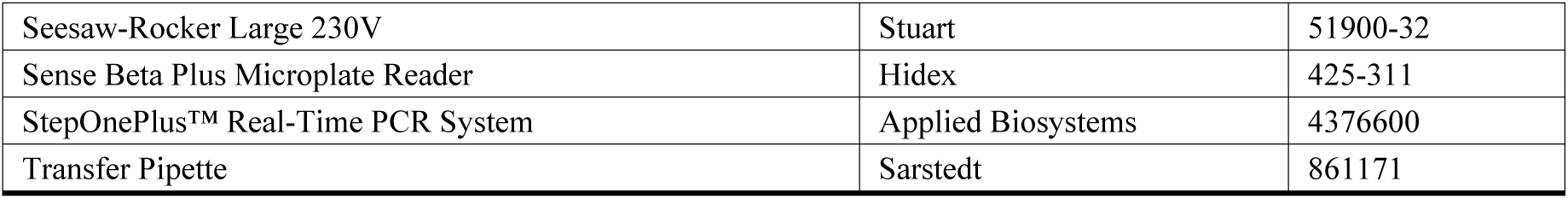

### RESOURCE AVAILABILITY

#### Lead contact

Further information and requests for resources and reagents should be directed to and will be fulfilled by the lead contact, Brian McDonagh.

#### Materials availability

All *C. elegans* strains generated in the laboratory will be available from the lead contact upon request.

#### Data and code availability

- Proteomic data has been deposited through ProteomeXchange (PXD059735) and the RNA-seq data has been deposited at GEO (GSE285634), they will be publicly available as of the date of publication. Accession numbers are listed in the key resources table. Original Western blot images and microscopy data reported in this paper have been deposited at Mendeley Data doi.10.17632/sztmnwvsd9.1.
- This paper does not report original code.
- Any additional information required to reanalyse the data reported in this paper is available from the lead contact upon request.

### EXPERIMENTAL MODEL AND SUBJECT DETAILS

#### C. elegans

*C. elegans* strains used in this study are listed in *KEY RESOURCES TABLE - Experimental models: Organisms/strains*. Nematode Growth Medium (NGM) was used to maintain worms.

#### Cell lines

C2C12 cells were grown in standard growth medium containing DMEM (Dulbecco’s Modified Eagles Medium, High glucose 4.5g/L), along with 10% foetal bovine serum (FBS) (Merck F7524) and 1% penicillin/streptomycin (P/S) at 37 °C in 5% CO_2._ The cells were regularly passaged every 2-3 days and maintained at 60%-70% confluence. For differentiation, cells were plated in 6 or 12 well plates. Cells were allowed to grow until cell density reached 80%, media was then changed to differentiation media, containing DMEM High glucose supplemented with 2% horse serum and 1% P/S (DM) and allowed differentiate for up to 7 days. Myotube area, diameter and fusion index were quantified as previously described ^60^.

### METHOD DETAILS

#### Tunicamycin (TM) treatment

TM treatment of C2C12 skeletal muscle myoblasts was performed when C2C12 myoblasts reached 70–80% confluency. A range of TM concentrations for 8h was used; after treatment, cells were either allowed to recover in DM overnight, myoblasts were prepared for analysis or differentiated in DM for 7 days.

TM treatment of *C. elegans* on a worm population synchronized by bleaching. Several plates of adult worms carrying embryos were subjected to a chemical (bleach-NaOH solution) and mechanical (by vortexing) procedure ^61^. The resulting embryos were treated with TM diluted in S Medium, for 24 h on a rocker at 20 °C. S Medium was obtained by centrifugation of an OP50 LB broth culture at max speed, the LB Broth supernatant was discarded and replaced with M9 Buffer. After treatment larvae were transferred to NGM plates and allowed to grow until adulthood.

#### Western blotting

C2C12 myoblasts were homogenised in a lysis buffer (150 mM NaCl, 20 mM Tris pH 7.5, 1 mM EDTA pH 8.3, 0.5% SDS, 1% Triton) [100 μL per well of a 6-well plate], on ice. Additionally, the lysis buffer was supplemented with Protease Inhibitor Cocktail [1 μL per 1000 μL of Buffer] and Phosphatase Inhibitor [1 μL per 1000 μL of Buffer]. Samples were prepared by scraping of cells using cell scrapers, followed by 30s homogenisation with an electric pestle. Protein extracts were quantified using the Bradford assay. Protein samples were diluted with Laemmli buffer (1 M Tris pH 6.8, 3% v/v glycerol, 2% v/v SDS, 1% v/v b-mercaptoethanol, 5 mg bromophenol blue and top-up to 10 ml H_2_O) to generate samples with the same protein concentration (1 μg/μL or 2 μg/μL). Unless otherwise specified, 20 μg of protein was loaded onto 12% SDS PAGE gels for Western blots. The proteins were transferred using a semi-dry blotter and the membrane was stained with Ponceau S to ensure complete transfer and for normalisation, membranes were blocked in 5% milk in TBS-T for 1 h at room temperature, when blotting for a phosphorylated protein 3% BSA in TBS-T was used. Subsequently, the membranes were incubated with primary antibodies (as listed in the materials table) at a 1:1000 dilution in 5% milk overnight, then incubated with secondary antibodies at a 1:10000 dilution in TBS-T in the dark for 1 h. Images were captured using the Odyssey Fc imaging system. Quantification and normalisation of the blots were performed using densitometry macros in ImageJ. Bands of the molecular weight of interest were selected (from the total protein or the Western blot membrane) and the density profile was plotted. Using the straight-line tool of ImageJ, a line was added at the base of the density profile, to remove the background signal. Finally, the area inside the profile was quantified. Normalisation was performed using total protein staining with Ponceau S. For two groups comparisons a two-tailed unpaired Student T-test was used, for more than two groups a one-way ANOVA was performed.

#### XBP1 PCR

After aspirating the culture medium from the plates, TRIZol was added directly onto the cells in each well: 0.3 ml for a 12-well plate. To ensure maximal RNA extraction, TRIZol was pipetted several times onto the well surface and then transferred into 1.5 ml Eppendorf tubes. For *C. elegans* samples age-synchronised worms from a single NGM plate were incubated with TRIZol (500 ml per NGM plate), samples were mixed by vortexing for 30s. Chloroform was added to each tube (100 μL for a 12-well plate and 1 ml for a *C. elegans* 60 mm Ø plate), and the tube was shaken vigorously by hand for 15s before incubation for 5min at room temperature. RNA precipitation was achieved by adding 100% isopropanol to the aqueous phase (250 μL for a 12-well plate) and incubating at -20°C overnight. RNA pellet was washed by adding 100 μL of 80% ethanol, supernatant was discarded and the tubes were allowed to air dry for 5-10min at room temperature. The RNA was reconstituted in 10 μL of RNase-free H_2_O. Samples were analysed using the NanoDrop2000 spectrophotometer to quantify nucleic acid content and purity. For the cDNA synthesis all samples were synthesised using 500 ng of RNA, ml random hexamers, 4 ml RT Buffer, 2 ml DTT, 1 ml dNTP, 1 ml Superscript II and 1 ml Ribolock. The samples were run in a thermocycler for 10min at 65 °C and 60 min at 42 °C; samples were diluted 10X (180 μL H_2_O) and kept at -20 °C until needed. For PCR a master mix was prepared containing: 1 μL of a primer mix Fwd+Rev (each 10 μM), 4 μL of H_2_O, and 6.25 μL of RedTaq for each sample. 12.25 μL of the master mix was added to each of the PCR tubes and then 1.25 μL of cDNA was added into each tube (total volume of 15 μL). Tubes were placed in a thermocycler with the following program: [*xbp-1u* and *xbp-1s*] 94 °C for 3 min, 94 °C for 30s, 58 °C for 30 s, 72 °C for 30s, 72 °C for 7 min (repeat steps 2 to 4 for 35 cycles), followed by 4 °C until analysis ^62^.

#### Silencing RNA and plasmid transfection

Transient knockdown of specific genes was performed using C2C12 myoblasts. Cells were plated on 12-well plates until they reached 70-80% confluency. Cells were treated with siRNA (50 nM) for 5 h in serum-free DMEM, using 4 μL/well of Lipofectamine 2000^TM^, for the SCR (non-transfected) group only Lipofectamine reagent was added. Media was refreshed with GM and cells were allowed to recover overnight before treatment with TM. Knockdown efficiency was validated by Western blotting of the proteins. Plasmid transfection was performed when C2C12 myoblasts reached 70-80% confluency on 24 well plates. For the transfection, 2 tubes were prepared in advance, each of which contained 125 μL/well of serum free DMEM. In the first tube, 2μL of lipofectamine was combined with the second tube containing 0.5 μg of plasmid and incubated for 5 min. The content of both tubes was combined by pipetting drop by drop the volume of the plasmid tube into the Lipofectamine tube and the final mix was incubated for 20 min. Finally, the transfection media was added to the cells drop by drop and they were incubated at 37 °C in 5% CO_2_ for 5h. The media was then refreshed with GM and cells were allowed to recover overnight.

#### Immunohistochemistry

Mitochondrial staining of myoblasts was performed following overnight recovery in DM after 8 h TM treatment. Medium was aspirated and cells were rinsed with PBS. Myoblasts were then exposed to mitochondrial dyes for content (MitoTracker green), and either membrane potential (TMRM) or ROS (MitoSox) diluted in serum-free DMEM for 20 min: MitoTracker Green 200 nM, TMRM 100 nM or MitoSOX 5 μM. In co-staining the dyes were incubated together. After a final PBS wash, images were captured using EVOS M7000. Regions within each cell showing staining fluorescence signals were identified as mitochondrial. For two groups comparisons a two-tailed unpaired Student T-test was used, for more than two groups a one-way ANOVA was performed.

*CTCF = Integrated Density – (Area of the cell X Mean fluorescence of background readings)*

MF20 immunostaining was performed after 7 days of differentiation. Samples were fixed using ice-cold methanol, after methanol was removed, samples were blocked with 10% horse serum in PBS for 1 h on a rocker at RT. Samples were incubated overnight with 1:500 primary MF20 antibody in 2% horse serum in PBS and 1:2000 of secondary antibody solution (anti-mouse 488) in 2% horse serum in PBS. Cells were incubated for 10 min in the dark at RT with 1:10,000 DAPI in PBS. Three parameters were quantified to assess the physiological adaptations of the myotube population. Myotube diameter was determined by measuring the thickest diameter of each myotube. The myotube area fraction, measures the percentage of the total image area that was occupied by myotubes. The fusion index, measures the fraction of the nuclei that are inside the myotubes and the total nuclei number, indicating their regenerative potential.

#### Viability Assay

The live/dead assay used a combination of ethidium bromide and acridine orange to distinguish between viable and nonviable cells. The protocol began with removing the culture medium and a rinse with phosphate-buffered saline (PBS). A staining solution was prepared by diluting ethidium bromide and acridine orange at 1:1000 in PBS, which was then added to the cells. The cells were incubated at room temperature on a rocker for five minutes, shielded from light by covering the plate. After incubation, the cells were imaged using an EVOS 7000 microscope, and the images were analysed with ImageJ software to quantify the number of live, dead, and necrotic cells. Each experimental group consisted of 3 replicates, with 4 images captured per replicate. Acridine orange stained all cells green, enabling the counting of both live and dead cells, while red-stained nuclei indicated cell death. Any red staining in the cytoplasm was considered either background noise or an indication of early necrosis.

#### Mitochondrial respiration rates

C2C12 myoblasts were plated in triplicate in XF HS Mini Analyzer plates at a density of 8000 cells per well in a total volume of 200 μL per well and incubated at 37 °C in 5% CO_2_ for 24 h. The cells were treated with TM for 8 h and the growth medium was refreshed with 10% growth medium to allow recovery overnight. Before analysis, cells were washed once with XF Real-Time Mito-Stress Assay Medium (containing 10 mM glucose, 1 mM sodium pyruvate, and 2 mM glutamine, pH 7.4) and the medium was replaced before incubating the cells at 37 °C in 0% CO_2_ for 1h. The concentrations of chemicals were optimised and based on the supplier instructions, 2 μM Oligomycin, 2 μM FCCP, and 1 μM antimycin A. Compounds were added to the hydrated cartridge, loaded with the plate and the calibration process was carried out in the XF HS Mini Analyzer.

For measuring mitochondrial respiration rates of *C. elegans*, the compounds (10 μM FCCP and 40 mM sodium azide) were added to the hydrated cartridge. Three well replicates were used that contained 8–13 worms per well and were transferred into an XF HS Mini Analyzer plate before starting the calibration process, all the procedures were performed at 20 °C ^63^. A one-way ANOVA analysis was used as the statistical test for group comparisons.

#### Mass spectrometry

Proteomics was performed to indicate the global label-free relative abundance of proteins. Protein extracts were using a hand homogeniser. Protein lysates were centrifuged at 15,000 g for 10 min at 4 °C and protein concentrations were determined using Bradford assay with BSA as the standard. Trypsin was reconstituted in 50 mM acetic acid, and 2 μg was added to the 100 μg of protein from samples, followed by overnight incubation at 37 °C. The digestion was stopped and RapiGest removed by acidification (3 μL of TFA and incubation at 37 °C for 45 min) and centrifugation (15,000 g for 15 min) ^64^.

Mass spectrometry analysis was performed at the proteomics facility NICB at Dublin City University. For LC-MS/MS analysis, an UltiMate 3000 RSLCnano LC system was coupled to an Orbitrap Fusion^TM^ mass spectrometer (Thermo Fisher Scientific). Peptides were loaded onto a trap column (AcclaimTM Pep-MapTM 100C18 LC Columns 5 μm, 20 mm length) for 3 min at a 10 μL/min flow rate in 0.1% formic acid (FA). Subsequently, peptides were transferred to an EASY-Spray PepMap RSLC C18 column (Thermo) (2 μm, 75 μm × 50 cm) operated at 45 °C and separated using a 60-minute gradient (buffer A: 0.1% FA; buffer B: 100% ACN, 0.1% FA) at a flow rate of 250 nL/min. The gradient ranged from 2% to 6% of buffer B in 2 min, from 6% to 33% B in 58 min, from 33% to 45% in 2 min, and held at 98% B for an additional 10 min. Peptides were ionised at 1.5 kV into the mass spectrometer via the EASY-Spray source, with a capillary temperature set to 300 °C. The mass spectrometer operated in a data-dependent mode, automatically switching between MS and MS/MS scans using a top 18 method (intensity threshold ≥3.6e5, dynamic exclusion of 20 seconds, and excluding charges +1 and > +6). MS spectra were acquired from 350 to 1400 m/z with a resolution of 60,000 FWHM (200 m/z). Peptide ions were isolated with a 1.0 Th window and fragmented using higher-energy collisional dissociation (HCD) with a normalised collision energy 29. MS/MS spectra resolution was set to 15,000 (200 m/z). The normalised AGC ion target values were 300% for MS (maximum ion time of 25 ms) and 100% for MS/MS (maximum ion time of 22 ms).

Mass spectrometry data was analysed for global changes in protein abundance. Raw files were processed using MaxQuant (v 1.6.0.16) against a *Mus musculus* protein database (UniProtKB/Swiss-Prot/TrEMBL, n sequences). Variable modifications included N-ethylmaleimide and D5 N-ethylmaleimide on Cysteine, oxidation of Methionine, and protein N-terminal acetylation. Minimal peptide length was set to 7 amino acids, and a maximum of two tryptic missed cleavages were allowed. Results were filtered at 0.01 false discovery rate (FDR) at both peptide and protein levels. The "proteinGroups.txt" file was analysed using Prostar software ^65^. Proteins with fewer than three valid values in at least one experimental condition were filtered out. A global normalisation of log2-transformed intensities across samples was performed using the local weighted regression (LOESS). Missing values were imputed using the structured least squares algorithm (SLSA) for partially observed values and DetQuantile for values missing from an entire condition. Differential analysis was conducted using the empirical Bayes statistics Limma. Proteins with a p-value < 0.01 and a Log2FC ratio > 0.58 or < -0.58 (FC > 1.5) were considered significantly changed, with an estimated FDR below 6.5% by Benjamini-Hochberg. For generating the volcano plots R 4.3.2 was used and the package ggplot2 3.4.4. Enrichment analysis and dot-plots were obtained with ShinyGO 0.80.

#### RNA sequencing and bioinformatic analysis

Total RNA was extracted from 4 individual NGM plates containing a population of *C. elegans.* The total RNA was extracted from an individual NGM plate for each condition following the TRIZol standard procedure described. A NanoDrop2000 was used to determine each sample’s purity and concentration. Samples were sent to Novogene for bulk RNA-sequencing on Illumina platforms. Novogene Sequencing Europe performed the initial bioinformatic analysis. The initial data analysis was performed by Novogene including sample and data quality control, alignment to the reference *C. elegans* genome using HISAT2 2.0.5, gene expression quantification with featureCounts v1.5.0-p3, differential expression analysis with DESeq2 v1.20.0 and gene set enrichment analysis with GSEA v4.3.2. The differential expression was obtained by normalisation of the raw readcount, the hypothesis test’s probability (p-value) was calculated by a statistical model and finally multiple hypothesis test corrections were used to obtain FDR values. The Multiple Testing online tool was used to calculate the FDR of the comparison ^66^. The Selected differential expression analyses were *EMBRYO CNT vs EMBRYO TM* and *L4 CNT vs L4 TM*. The screening criteria for differential genes used were |log2 (FoldChange)| >= 1 & padj<= 0.05. R 4.3.2 was used for the volcano plots and the package ggplot2 3.4.4. Enrichment analysis and dot-plots were obtained with ShinyGO 0.80. Pathway analysis of RNA seq. Data was visualised using PathVisio 3.3.0 to highlight altered pathways from detected transcripts.

#### Ca^2+^ Imaging

Myoblasts were plated on 12-well plates. After recovery overnight in DM following the TM treatment, cells were washed 2 times with Ca^2+^-containing medium (NaCl 135 mM, KCl 5 mM, MgCl_2_ 1 mM, CaCl_2_ 2 mM, HEPES,10 mM and Glucose 10 mM, pH 7.45). Cells were loaded with 2 μM Fluo-4/AM (5 μM) for 40 min in a 37 °C incubator in the dark. The cells were washed twice in Ca^2+^-containing medium and left in the dark for an additional 15 min to allow for the de-esterification of Fluo-4/AM. Media was replaced with ∼1 ml of Ca^2+^-containing medium, images were recorded using the EVOS M7000. Image sequences were collected over a 20min video at 0.5 Hz. To assess Store-Operated-Ca^2+^-Entry (SOCE), which is an indicator of Ca^2+^ level in the endoplasmic reticulum, the classical thapsigargin/Ca^2+^ re-addition protocol was followed: the cells were left for 5 min in Ca^2+^-containing medium. The media was replaced with Ca^2+^-free medium (NaCl 135 mM, KCl 5 mM, MgCl_2_ 1 mM, EGTA 1 mM, HEPES 10 mM and Glucose 10 mM, pH 7.45). After 2 min 1 μM Thapsigargin (irreversible blocker of SERCA pump activity) was added to deplete the internal Ca^2+^ stores. After 8 min, rapidly replace the medium with the Ca^2^-containing medium (Ca^2+^ will enter the cells through the SOCE channels). Following 5 min the timelapse was terminated. Timelapse videos were processed and analysed using FluoroSNAP in MATLAB (MathWorks, Inc.). Myoblasts exhibiting fluorescence changes > 5% during recording were identified through time-lapse analysis, and cell boundaries were determined via batch segmentation. A time-varying fluorescence signal was then calculated and expressed at ΔF/F0, a threshold of 0.05 was applied to identify events. The frequency, amplitude, duration, and synchronization of spontaneous and evoked Ca^2+^ transients were analysed using a custom script in the R software ^67^. Student T-test was used to compare the two groups.

#### Transmission Electron Microscopy

C2C12 myoblasts, myotubes and *C. elegans* were fixed using 2% glutaraldehyde and 2% paraformaldehyde in 0.1 M sodium cacodylate buffer at pH 7.2. Following fixation, the buffer was replaced with 1% osmium tetroxide in 0.1 M sodium cacodylate buffer (pH 7.2). Subsequently, samples underwent dehydration using a series of ethanol concentrations and were infiltrated with resin-acetone mixtures overnight, starting with a 50:50 mixture, a 75:25 mixture, and finally 100% resin. Ultra-thin sections (70–90 nm) were cut in a sagittal plane using a diamond knife and mounted onto 3 mm copper grids. The grids were then counterstained for 5 min with 2% uranyl acetate and 5 min with 2% lead citrate, after that they were examined under a Hitachi 7500 TEM microscope at a magnification of 10000x, 25000× and 60000x. Traditional point-counting methods assessed cell and organelle composition ^68^. A student T-test or one-way ANOVA was used to compare the groups. The first parameter calculated, in the X10000 augment’s images, was the volume fraction (V_V_) of mitochondria (an indicator of mitochondrial content), a 0.25 mm grid was used and the grid intersections with the organelles and tissues were counted and the following formula was applied ^68^:

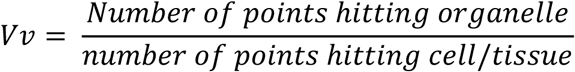

The X25000 augment’s images were used for the calculation of the mitochondrial aspect ratio (indicator of mitochondrial elongation), in this case the major and minor axis were measured and the following formula was used ^68^:

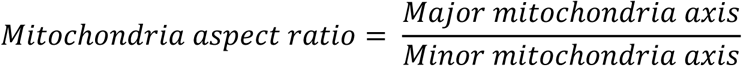

Then X60000 images were used for calculating the surface density (Sv) of membranes (in general indicators of abundance), test lines were superposed to our images (length = 2156 nm, separation = 100 nm) and the intersections with membranes were counted and used in the following formula ^68^:

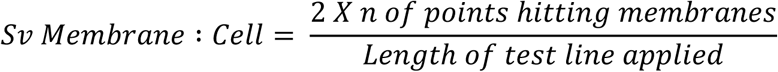

The relative estimates were combined with nuclear volumes to provide estimates of cell and organelle volumes and surface areas as follows. The 4-way nucleator method was applied to the profile to study the nuclear volume, and the distance from the nucleolus to the nuclear membrane was measured and averaged (Ln) ^68^.

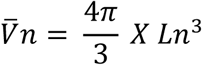

To estimate the cellular volume, the volume fraction (V_V_) of the nucleus was calculated ^68^:

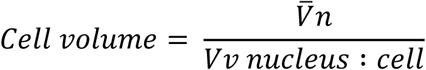

The membrane surfaces were calculated using the formula ^68^:

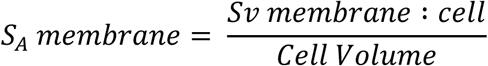

#### Lifespan assays

One hundred L4 worms were transferred to NGM plates supplemented with 5-fluorouracil 10 mM and incubated at 20 °C. 10 μM 5-fluorouracil (5-FU) was added to the NGM plates. Worms were then allowed to grow in normal NGM plates until L4 stage and transferred to the 5-FU supplemented NGM plates. Lifespan assay was initiated on day1 of adulthood and their viability was scored every two days. Worms were considered dead when the animal did not respond to gentle touch with a pick, animals were censored when they died by desiccation, ruptured vulva or internal hatching. Data was analysed using Kaplan–Meier survival analysis with the Log-rank test, when comparing two conditions. For multiple comparison survival analysis, χ^2^ and Bonferroni corrected p-value (BPV) were calculated for each condition, using the online platform for statistical analysis of survival data, OASIS 2 ^69^.

#### Oxidative stress survival assays

On day 1 of adulthood 50 worms were transferred to a 24-well plate, each well containing 250 μL of M9 buffer. Each test well contained had a final concentration of 10 mM NaAsO_2_ or 100 mM of Paraquat in M9 buffer. Viability was scored every hour and the worms were considered dead when the animal did not respond to gentle touch with a pick ^70^. For the tBuOOH stress assay NGM plates containing 15.4 mM tBuOOH were used. 50 worms were transferred to the test plates along with some OP50 to avoid their escape from the plate (during the first hour of the assay the worms were monitored every 10 min to avoid desiccation when trying to escape the plate). Viability was scored every hour and the worms were considered dead when the animal did not respond to gentle touch with a pick ^71^. When comparing two conditions, data was analysed using Kaplan–Meier survival analysis based on Log-rank test. For multiple comparison survival analysis, c^2^ and BPV were calculated for each condition, using the online platform for statistical analysis of survival data, OASIS 2 ^69^.

#### Microscopy of *C. elegans*

MitoTracker and MitoSOX staining on *C. elegans* were conducted on 45 worms for each set of conditions. The worms were incubated with either 50 μL of 2.5 μM MitoTracker Red CMXRos or 10 μM MitoSOX Red, diluted in M9 buffer, for 10 min or 1 h, respectively, at 20 °C in darkness. Worms were transferred to seeded NGM plates and allowed to forage in darkness for 2 h at 20 °C to prevent stain accumulation in the guts. Subsequently, 15 worms per replicate, for a total of 3 replicates were immobilised on unseeded NGM plates using a drop of 20 mM Levamisole. All steps were conducted in darkness. Imaging was performed using EVOS at 10X magnification, and ImageJ was utilised to evaluate the CTCF of each worm. 15 worms per replicate were imaged, for a total of 3 replicates, that were immobilised on unseeded NGM plates and imaged by EVOS at 10X magnification. ImageJ was used to assess the GFP CTCF for each worm. In all strains that required length measurements, 15 worms per replicate, for a total of 3 replicates were immobilised on unseeded NGM plates and imaged by EVOS at 10X magnification. The length measurements were assessed using the *segmented line* tool from ImageJ.

LysoSensor green and LysoTracker staining were conducted on *C. elegans* by soaking in 200 mL M9 buffer containing LysoSensor Green DND 189 and LysoTracker red DND 99 10 mM for 1 h at 20 °C in the dark. Worms were then transferred to fresh NGM plates with OP50 and allowed to recover at 20 °C for 1 h in the dark. Imaging was conducted at 60X magnification with an EVOS M7000 microscope and the LSG/LTR fluorescence in the intestine area for each worm was then analysed using ImageJ.

Oil red O staining was conducted as previously reported ^72^. Animals were mounted on glass slides and imaged with the EVOS M7000 microscope colour camera. Oil red O was quantified from colour images using the level of excess red intensity in the red channel compared to the blue and green channels as described ^31^.

The physiological stress reporters *hsp-4p::GFP, hsp-6p::GFP and gst-4p::GFP* were imaged using the standard procedure adapted from above. Prior to the imaging an induction of ER or mitochondrial stress was performed. *hsp-4p::GFP* animals were grown on NGM from hatching until L4 at 20 °C, treated with ER stressor 25 ng/ μL TM on supplemented OP50 plates for 16 h at 20 °C. *hsp-6p::GFP* animals were grown on NGM from hatching until adult day 1 at 20 °C, they were then treated with the mitochondrial uncoupler 50 μM CCCP on supplemented OP50 plates for 8 h at 20 °C prior to imaging. *gst-4p::GFP* animals were grown on NGM from hatch until adult day 1 at 20 °C and were treated with 50 µM Paraquat in 2 ml M9 Buffer for 2 h at 20 °C, followed by a recovery in NGM plates for 2 h at 20 °C.

The *myo-3::mitoGFP* reporter strain for muscle mitochondrial morphology was imaged using the EVOS M7000 microscope at 60X magnification. On day 1 of adulthood, 45 worms per condition were immobilised with 20 mM Levamisole on a 4% agar pad slide and covered with a coverslip, sealed with nail polish to allow visualisation with immersion oil. Mitochondrial morphology between the pharynx and vulva was evaluated based on three classifications: filamentous (showing filamentous mitochondrial networks), punctate (displaying mostly fragmented mitochondria), and intermediate (exhibiting isolated mitochondrial networks) ^73^. The reporter strain used to monitor mitophagy was *p_myo-3_TOMM-20::Rosella* (a gift from the Tavernarakis lab). The standard procedure described above was used, 15 worms per replicate, for a total of 3 replicates were immobilised per replicate onto NGM plates. Imaging was conducted at 40X magnification with an EVOS M7000 microscope and the CTCF of green and red fluorescence in the head area for each worm was then analysed using ImageJ, to obtain the GFP/DsRed ratio.

#### CeleST Swimming activity assay

Swimming behaviour of the different *C. elegans* strains was assessed at day 1 of adulthood or as indicated. On the day of the assay, a glass slide with a 10 mm pre-printed ring was prepared and loaded with 50 μL M9 Buffer. Five worms were picked and placed into the swimming area, for 45 animals per condition. When worms were transferred to liquid media they started swimming very quickly, to avoid false activity recordings they were allowed to rest in the M9 Buffer for 20 sec before starting the recording. 30s movies with ∼16 frames per second of the animals were taken using a Nikon LV-TV microscope at 1X magnification with a OPTIKA C– P20CM camera. These videos were exported to and processed by the CeleST software ^29^. Student T-test or one-way ANOVA was used to compare the groups.

### QUANTIFICATION AND STATISTICAL ANALYSIS

Statistical data were represented as mean ± SEM. For comparisons of means between two groups a two-tailed unpaired Student T-test was performed (data normality was validated using GraphPad prism). For comparisons involving more than two groups, one-way or two-way analysis of variance (ANOVA) was used. Lifespan and survival assay data were analysed using Kaplan–Meier survival analysis with a Log-rank test when comparing two conditions. For multiple comparison survival analysis, χ^2^ and Bonferroni corrected p-value were calculated for each condition, using the online platform for statistical analysis of survival data, OASIS 2 ^69^. chi-square test was used to compare the distribution into multiple categories. Prism version 9.5 software package for MacOS was used for the statistical analysis. A p-value < 0.05 was considered statistically significant and the significance indicators were assigned: * p ≤ 0.05, ** p ≤ 0.01, *** p ≤ 0.001 & **** p ≤ 0.0001.

The following procedures were employed for the different data analyses and quantification. Images obtained from Western blots and microscopy of *C. elegans* were subjected to semiautomated quantification using ImageJ and manual corrections were applied to ensure accuracy during the quantification process. Western blot band intensities were normalised according to total protein intensity from Ponceau S staining. For C2C12 microscopy, at least 3–6 images were captured randomly at 10X, 40X or 60X magnification from different fields of view per each biological replicate. MF20 immunostaining myogenic potential analyses, in each field of view, average diameter of all myotubes was measured, and the average area fraction of myotubes and average fusion index were determined using the percentage of nuclei within myotubes against the total number of nuclei. For fluorescence intensity, the relative fluorescence intensity per cell/worm was assessed and corrected with cell area and background signal. For the microscopy of *C. elegans*, at least 45 worms were measured at 10X or 60X magnification per condition. For the SJ4103 strain, images from body wall muscle mitochondria network morphology from the area between the animal’s pharynx and vulva were blindly distributed in three categories: filamentous, punctate and intermediate. For the mitochondrial dynamics, the assessment involved determining of each worm’s ratio of green fluorescence to red fluorescence.

